# Stability of multi-species consortia during microbial metabolic evolution

**DOI:** 10.1101/2025.01.15.633269

**Authors:** Dan Kehila, Alireza G Tafreshi, Nobuhiko Tokuriki

**Affiliations:** Michael Smith Laboratories, University of British Columbia, 2185 East Mall, Vancouver, V6T 1Z4, BC, Canada; Biodiversity Research Centre, University of British Columbia, 2212 Main Mall, Vancouver, V6T 1Z4, BC, Canada

## Abstract

Explaining multi-genic adaptations is a major objective of evolutionary theory. Metabolic pathways require multiple functional enzymes to generate a phenotype, and their evolution in microbes remains underexplored. In particular, sites polluted with manmade chemicals or “xenobiotics”, like plastic or pesticides, provide evidence for the rapid adaptation of novel metabolic pathways in microbes, which degrade these xenobiotics into utilizable nutrients. Decades of microbiological studies revealed that these pathways often are not consolidated within a single microbial species, but are rather distributed across several different ones, which cooperatively degrade xenobiotics. These species form remarkably stable consortia in the laboratory, but the determinants of this stability have not been hereto addressed. In this study, we show that trade-offs in microbial life history explain stable co-existence in a mathematical model of a three-species consortium, growing on a xenobiotic as the sole source of a limiting nutrient. Stability is predicted to hinge on a specific “ecological matching” between a species’ metabolic role in the novel metabolic pathway and its nutrient utilization strategy.

## Introduction

Catabolic pathways comprise a series of chemical reactions, each catalyzed by specific enzymes, that collectively convert substrates into usable nutrients. The evolution of metabolic pathways is complex, requiring the simultaneous presence of multiple genes, each encoding a necessary pathway enzyme, in order to display a selectable phenotype^1– 3^. Possessing only a subset of the pathway genes results in incomplete catabolism, no nutrient production, and therefore no fitness advantage^4–6^. While theories explaining the evolution of a single enzyme-catalyzed reaction are well-established, primarily through the latent “promiscuous” activities of existing enzymes, which is subsequent refined by natural selection^7–11^, applying this model to entire metabolic pathways is challenging. This is because the probability of sourcing multiple promiscuous enzymes within a single microbial population decreases geometrically with the length of the pathway. It is this explanatory challenge that has prompted much research into other evolutionary models that can plausibly generate an ancestral pathway from standing genetic variation^1,8,12,13^

Recently discovered manmade chemical (xenobiotic) catabolic pathways, such as those for atrazine and *penta*-chlorophenol, suggest that these pathways evolved through the recruitment of multiple enzymes from different organisms via horizontal gene transfers (HGTs)^6,14,15^. These observations led to a hypothesis that novel, multi-enzyme pathways originate from promiscuous enzymes distributed across various species within a microbial community^6,12,16^. This hypothesis originated from research on Enhanced Biodegradative Consortia (EBC)^6,17,18^: groups of microbes routinely enriched by microbiologists trying to isolate xenobiotic-degrading microbes in polluted sites. In those studies, a single microbial isolate, which can degrade xenobiotics independently, was rarely found. Instead, EBC were enriched, where each consortium member catalyzes specific step(s) of the pathway (intermediate metabolites being transferred between members) and the consortia collectively degrade the xenobiotic. This community-based metabolism results in a unique, sequential cross-feeding scenario that benefits all members^6,12,17,18^. Given the genetic and metabolic diversity of microbial communities, the formation of stable interspecific relationships evident in EBC supports the rapid evolution of catabolic pathways in a community context, which can be followed by enzyme optimization, gene duplication and HGT to consolidate these enzymes into a complete pathway in a single organism.

While community-based metabolism helps explain the rapid evolution of new catabolic pathways^12^, it raises new questions about ecological stability^6^. All members that contribute to the pathway must stably co-exist within the consortia to sustain a metabolic flux. If even a single species grows too slowly and is diluted to extinction, this could lead to the diminishment or disappearance of a biochemical step, and the entire pathway and consortia can collapse. Despite this potential vulnerability, EBC often exhibits exceedingly stable compositions, with all metabolizing species maintained for months or years while growing on the xenobiotic as the sole source of an essential nutrient^18^. Currently, the question of ecological stability remains unanswered.

The purpose of this article is to study a model for the stable co-existence of a microbial consortium that performs community-based metabolism, such as the EBC described above. We highlight two pertinent questions regarding the stability of EBC, arising from the metabolic “roles” different species find themselves in at the early stage of the nascent metabolic pathway, and propose answers using our model. Exploring the coevolution of this stable-coexistence after group formation is beyond our scope, instead we focus on demonstrating that, under laboratory conditions, metabolic differences between species are sufficient, and arguably necessary, to generate stable coexistence in community-based pathways.

## The model

### A model of consortium-based sequential metabolism

We develop a simple model for a microbial consortium containing a novel metabolic pathway distributed across different species, considering three representative species “types” that could be found in an EBC (**Figure 1**). “*I* species” possess enzyme(s) that perform intermediate reaction(s), whereas “*T* species” has the enzyme required to conduct the terminal, nutrient-generating reaction. A terminal reaction can be thought of as producing a metabolite that is no longer foreign to the microbes and pre-existing in its biological network (and thus usable as a nutrient source). Finally, a free-riding or “*F* species” does not play a metabolic role in the pathway but remains within the enrichment, thus existing as a “free-rider” on nutrients liberated from *T*. In fact, since the *I* species does not produce a chemical that is directly usable as a nutrient (only the intermediate), it too depends on *T* for nutrient benefits^6^.

**Figure 1.**
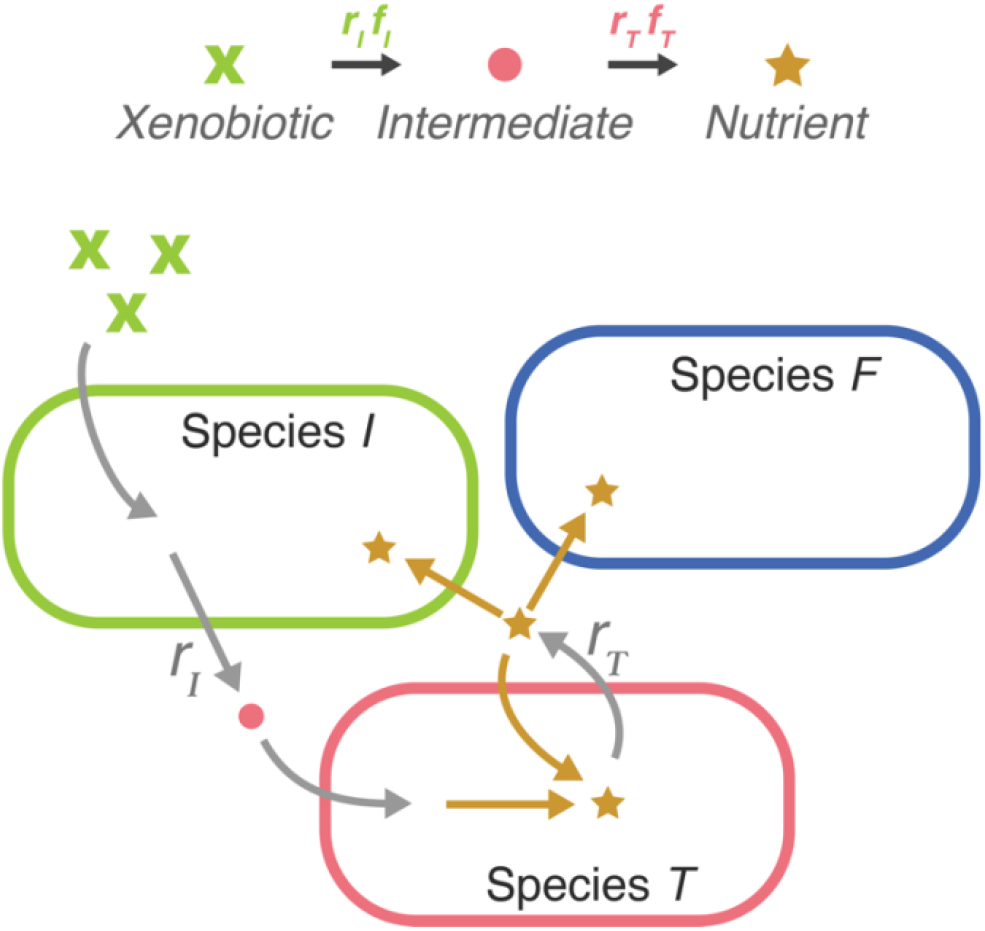
Cartoon of a three-species consortium growing on a xenobiotic (green X) as a sole nutrient source. Species *I* converts the xenobiotic into an intermediate (red dot), which it cannot consume, at a rate *r*_*I*_ . Species *T* is metabolically capable of liberating a usable nutrient (yellow star) from the intermediate. The growth of *I* and species *F*, which does not participate in the xenobiotic biodegradation pathway, depends on the rate *r*_*T*_ at which *T* generates a publicly available nutrient pool from its growth on the intermediate. Conversion rates are also a function of the frequency of the converting microbe in the consortium (see text).

Our model is designed to address two challenges arising in the stable co-existence of EBC:

1. *Asymmetric benefits to mutualists* (Section 1): though the two metabolizing species, *I* and *T*, engage in a mutualistic cross-feeding relationship, a potential asymmetry of benefits, e.g., accessibility to the nutrients, exists between them. Namely, *T* species generate nutrients and thus presumably have some preferential access to them. In contrast, *I* require access to the nutrients liberated by *T*, typically assumed to arise through passive leakage or cell lysis^17,18^ (yellow arrows in **Figure 1**). Clearly the asymmetry favors *T*, and possibly endangers the other species (*I*, in this case) and the viability of the consortium. Thus, remarkably, we expect the stability of an EBC composed only of mutualists to still be compromised even in the absence of free-riding. Since EBC are enriched on a xenobiotic as a sole nutrient source, asymmetric benefits may be a common feature, expected whenever non-nutrient generating species exist.
2. *Well-mixed environment* (Section 2): Standard EBC enrichments utilize chemostats^12^, specialized bioreactors that maintain constant mixing, inflow and outflow rates during culturing, ensuring well-mixing, constant xenobiotic concentrations, and culture growth rates^19^. Well-mixed environments are often regarded as discouraging, if not inhibiting, ecological co-existence^20–23^ or the cooperative sharing of nutrients between microbes as “public goods”^24,25^ (but see^26^). In EBC enrichments, well-mixing ensures that both non-terminal reaction species (*I* and *F*) have equal access to a public pool of nutrients liberated by *T*. As this public pool is the sole and limiting nutrient source for *I* and *F*, they are perfect competitors, making their co-existence theoretically unstable. The prevalence of stable EBC, containing multiple species with equal access to a single limiting nutrient, must be reconciled with theory.

Stability within our model EBC requires some constraints on its members to offset asymmetric benefits and well-mixing. We hypothesize these constraints arise from the differential nutrient utilization strategies of each species, which a central feature of our model and discussion. Microbes show specialization in their ability to uptake and utilize nutrients, based on environmental nutrient concentrations^36–38^. Given that nutrient concentrations, in turn, correlate with the density of metabolizing species in the culture, evolutionary game theory serves as a suitable framework for our analysis^27–30^. Specifically, we model the growth rate of species *j* as a payoff ***P***_***j***_, which correlates linearly with the concentration *C*_*j*_ of all nutrients it can consume, such that

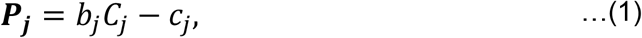

where *b* and *c* are species-specific parameters, representing a species’ ability to convert nutrients into reproductive benefits, and to survive in the absence of nutrients, respectively. These roughly correspond to the two growth regimes mentioned above: when nutrients are plentiful, payoffs are mainly determined by the *b* parameter, or otherwise by *c* when nutrients are scarce. Note that a biological definition of the parameters *b* and c requires some care. Real microbial nutrient-growth relationships are seldom, if ever, linear^31^, making this payoff approach linear regression-like. It is therefore important to restrict the interpretation of the parameters as useful approximations of growth when nutrients are plentiful vs. scarce. With this in mind, *b* should be constrained to the positive reals, but *c* is most flexibly employed when allowed to take either positive or negative values.

The definition of these growth parameters also lends itself to a possibility for an elementary analysis of microbial life history. It is well-known from as early as the work of Monod and colleagues that a microbe’s ability to survive and grow on scarce nutrients trades off with its growth rates when nutrients are plentiful^31–35^. Here, we utilize *b* and *c* to analyze distinct strategic classes of growth, known as oligotrophy and copiotrophy^36– 38^. Briefly, these respectively correspond to high survival in scarce nutrient and slow growth on ample nutrients (small *b* and low *c)*, and conversely, poor survival in scarce nutrients and rapid growth rates on ample nutrients (large *b* and a high *c)*. We explore the consequences of these trade-offs on co-existence in our model.

To model asymmetric benefits, we assume *I* and *F* species can only utilize the nutrients that are generated by *T*, in proportion to its concentration *C*_*NUT*_; thus in their respective payoffs we have *C*_*I*_ =*C*_*F*_ =*C*_*NUT*_. In contrast, *T* can utilize both nutrients and intermediates to grow, such that *C*_*T*_ =*C*_*NUT*+_*C*_*INT*_. We assume each metabolizing species *j* performs a metabolic step at a fixed rate *r*_*j*_, representing the rate at which the product of *j*’s metabolism is made available in the culture by *j*.

Assuming well-mixed conditions, we represent the concentration of *C*_*INT*_ as directly proportional to *I*’s frequency in the enrichment; specifically, it is the product of the frequency of species *I* times its conversion rate, *f*_*I*_ *r*_*I*_. Modelling *C*_*NUT*_ is more complex, however, since it is generated by a flux through the pathway carried out by *I* and *T*, rather than a single reaction. To address how this flux depends on the relative frequencies of the metabolizing species, we explore two possible definitions. In the first, *C*_*NUT*_ is defined as the product of the rates of both steps required in their production; more generally, one would represent the concentration of metabolite *s* in the pathway as 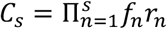. This representation requires that both species are present for nutrient production, but unrealistically assumes that nutrient generation is highest when the two species involved in its production are exactly balanced (*f*_*I*_ =*f*_*T*_). In a second definition, which is presented in Appendix A, we use a more realistic representation for nutrient generation based on metabolic control theory^39^; this relaxes the above assumption, though the qualitative outcomes presented here remain unchanged.

The determinants of stable co-existence are studied by obtaining solutions to the system of differential equations

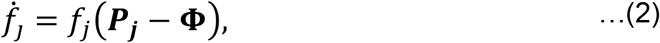

where **Φ** = Σ_j_[***P***_***j***_*f*_*j*_] is the average fitness. Solutions to (2) yield fixed points, and those fixed points that are stable and contain all consortium members are analyzed. Deterministic equations and continuous changes in frequencies assume large population numbers, especially for dynamics at the limit of very small frequencies. This is appropriate for the typical conditions of EBC enrichments, where culturing volumes (volume of vessel) is usually on the order of several liters, corresponding to microbial densities on orders exceeding 10^12^. Chemostat conditions justify our assumption of well-mixing, and also of using frequencies as proxies for concentrations, as xenobiotic concentrations themselves are not limiting (though nutrient production is still limited by the flux through the consortium-based pathway). Finally, note that with a large population, changes in frequency are gradual enough that a linear response in concentrations with frequency change is suitable. More delayed or rapid changes in concentration, or growth rates, from small changes in frequency may result in oscillatory dynamics, which are outside the scope of this study.

## Results

### Section 1: Stable co-existence of *I* and *T* mutualists

First, we explore the effect of asymmetric benefits between *I* and *T* species in the absence of *F*. Their respective payoffs are

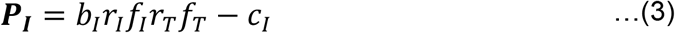

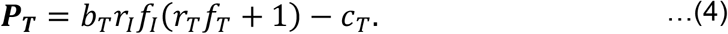

Clearly, the payoff differences ***P***_***I***_ **− *P***_***T***_ given *b*_*I*_*=b*_*T*_ and *c*_*I*_*=c*_*T*_ is always negative, demonstrating our expectation for asymmetry between the two species. Accordingly, the mutualist pair can stably co-existence if species *I* possesses some form of physiological advantage relative to *T*; for instance, *I*’s ability to convert nutrients into growth better than *T* (*b*_*I*_ *> b*_*T*_) or surviving better when nutrients are scarce (*c*_*I*_ < *c*_*T*_). The parameter space enabling stable co-existence, when *I* has a physiological advantage, is shown in the white region of **Figure 2A**, using parameters 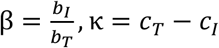 to capture the differences in growth parameters between *I* and *T*. The choice of a ratio for *β* but a difference for *κ* stems from the aforementioned consideration of the parameter domains. Evidently, *b* or *c* alone is sufficient for stable co-existence of both species (see Supplementary Information).

**Figure 2.**
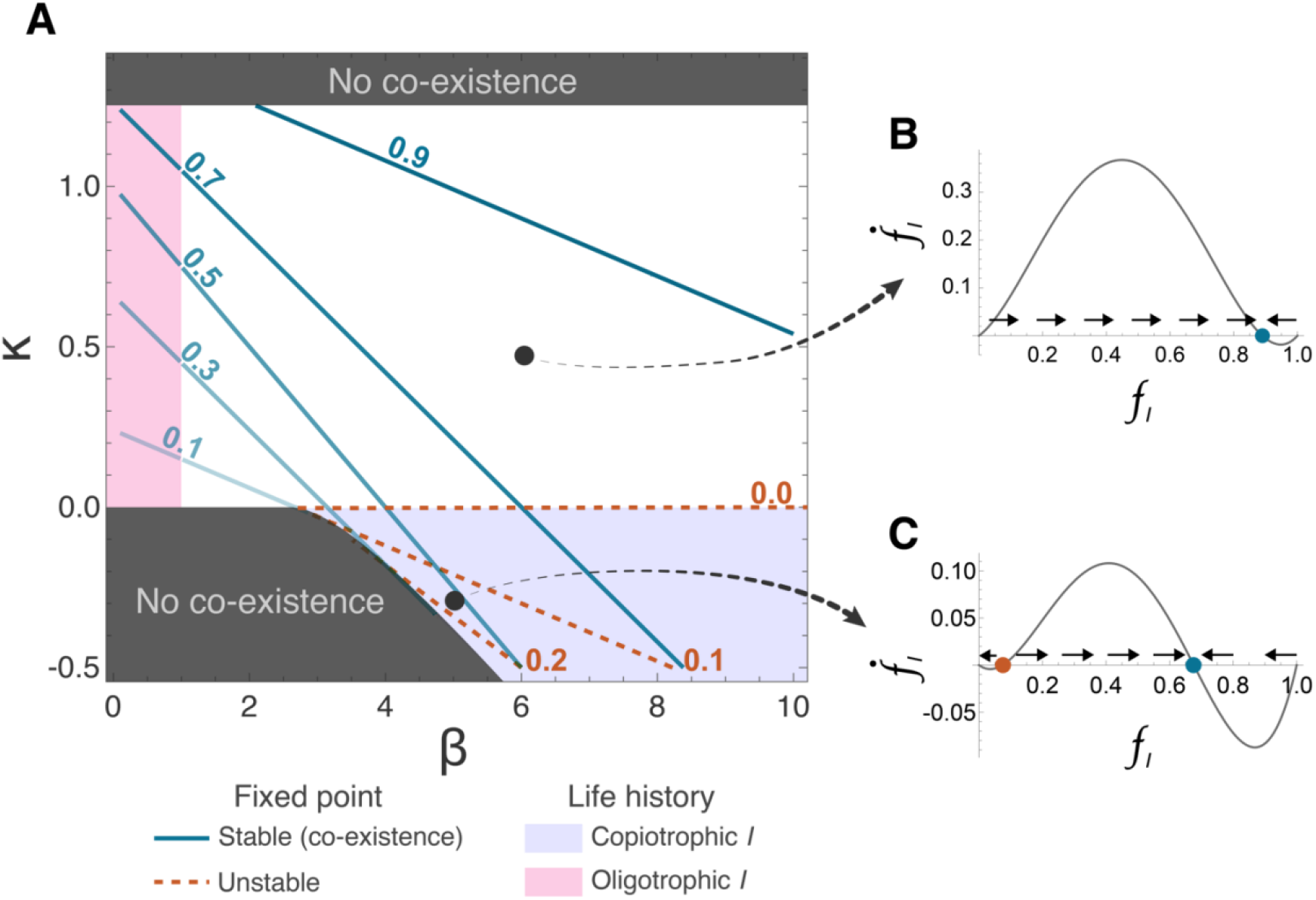
(A) Contour plot showing how advantages in the payoff parameters impact the equilibrium fraction *f*_*I*_ in the *I,T* consortium. Blue lines indicate stable *f*_*I*_ equilibrium frequencies (fixed points), and red lines indicate unstable equilibrium (below which *I* decrease to extinction). Upper and lower bounds are shown as dark grey lines. Parameter regimes for copiotrophy and oligotrophy are filled in. Additional parameter values used were *r*_*T*_ = 1, *b*_T_ = 1, *r*_*I*_ = 5/4. (B) Example of *I,T* consortium dynamics (change of *I* frequency for any given frequency *f*_*I*_) showing one stable equilibrium (blue dot), evaluated at *κ* = 0.5, *β* = 6. (C) Example of *I,T* consortium dynamics with copiotrophic *I*, showing one stable equilibrium (blue dot) and one unstable equilibrium (red dot), evaluated at *κ* = **−**0.25, *β* = 5.

The parameter space in **Figure 2** further reveals that stable co-existence is possible even when *I* shows a disadvantage in one of the growth parameters, provided that it has an advantage in the other. Thus *I* need not necessarily display physiological dominance, but can show a relative physiological specialization to *T* as well. Physiological specialization can arise if the two species adopt distinct life history strategies—e.g., *I* growing better on ample nutrients, but worse on scarce nutrients, than *T*. Specifically, two parameter regimes (*β>1* and *κ < 0)* or *(β <1* and *κ* >*0)* respectively represent a copiotrophic or oligotrophic *I*, relative to *T*, where *I* has an advantage in one parameter and disadvantage in the other, and co-existence nevertheless remains stable. These regimes correspond to the colored regions **Figure 2**. For the remainder of the section, we analyze the determinants of stable co-existence within these two specialization regimes.

C*opiotrophic I species (β>1 and κ < 0; purple shaded region in* ***Figure 2***): in this regime, the interaction exhibits two fixed points, one stable, and the other unstable for lower values of *f*_*I*_. The unstable fixed point (depicted by red dashed contours in **Figure 2**) arises because a copiotrophic *I* relies on ample nutrients to gain its advantage over *T*. However, when *I* is rare, *T* obtains few intermediates and the public pool of liberated nutrients remains small. Thus a rare *I* faces a hurdle in avoiding its own extinction when nutrient liberation by *T* is too slow. This dynamic leads to the unstable equilibrium, requiring *I* to exceed a certain threshold frequency for the consortium to reach stable species co-existence (**Figure 2c**).

Studying the copiotrophic parameter regime that allows co-existence of both species, we first recognize that *I*’s growth advantage must compensate for deficiencies in *T*’s public nutrient production rate *r such* that 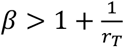. If *r* is too low, nutrients are not sufficiently available for *I* to obtain an advantage. Additionally, stable co-existence requires the trade-off between *β* and *κ* to satisfy

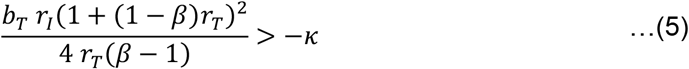

Note that both copiotrophy conditions imply advantages in the *b* parameter are unbounded. Equation (5) specifically shows that *β* and *κ* can diverge to arbitrary magnitudes, in line with the well-known characteristic of copiotrophs displaying vast variations under nutrient-rich versus nutrient-poor conditions (feast vs. famine)^38^.

*Oligotrophic I species (β <1 and κ >0; pink shaded region in* ***Figure 2***): in this regime, we first establish that *κ* must not be so high as to constitute the bulk of *I*’s payoff. Otherwise, *I* becomes unrealistically self-sufficient, and the system ceases to represent a co-dependent consortium. We thus expect the oligotrophic advantage to be bounded. In our model, this boundary is defined where *I’s* baseline payoff *-c*_*I*_ is lower than the maximum payoff *T* can attain when only intermediates are available (as *f*_*I*_ approaches 1). In other words, we require that

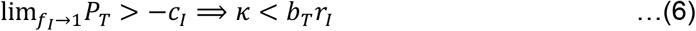

Beyond this upper bound for *κ, I* outgrows *T* regardless of xenobiotic catabolism, and it would be unusual to classify *I* as “oligotrophic”. Consistent with our expectations, *κ* is bounded both from below at 0, and above by *T*’s growth rate on the intermediate, unlike the unbounded advantage term in the case of copiotrophy.

### Section 2: Stable co-existence of *I* and *T* with a free-riding species *F*

Next, we consider the interactions among three species; *I* and *T* populations involved in xenobiotic degradation, and an additional non-metabolizing *F* species. We assume a well-mixed environment, meaning nutrients released by *T* are equally available to *I* and *F*. Thus both species experience *C*_*NUT*_ in the same manner, though the *b* and *c* parameters could still vary in a species-specific manner, as described above. In the absence of spatial structuring, previous studies of co-existence driven by negative frequency-dependence often rely on non-linear payoffs to explain stable co-existence of multiple species or genotypes, incorporating assumptions such as saturating payoffs or a threshold frequency of nutrient producers necessary for feeding others^29,30^. However, even if we incorporate such features into equation (1), making payoffs some nonlinear function *Ψ*(C_*NUT*_) of nutrient concentrations, the functional form for both *I* and *F* must remain the same, as they both experience the same nutrient environment. Without species-specific differences, the frequency-dependent terms will cancel out in the payoff difference ***P***_***I***_ *–* ***P***_***F***_ regardless of whether *Ψ* has non-linear features. Conversely, if the parameters in the function *Ψ* are allowed to vary between *I* and *F*, as we will show, a linear form (1) is sufficient to describe stable co-existence. Our assumption of species-specific parameters, allowing for differential nutrient utilization and life history specialization on the same nutrient source, is therefore not only sufficient but possibly also necessary to explain stable co-existence in well-mixed EBC enrichment conditions.

A three-membered extension of the payoff system described in equations (3) and (4) now includes the *F* species, where

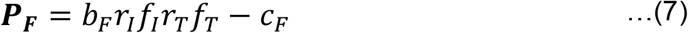

For this system of three equations there exists a single fixed point containing all three species on a single limiting nutrient; this fixed point reveals an interesting emerging pattern, depicted in **Figure 3**. Recall firstly, that in the case of two species (*I* and *T* above), co-existence required variation in only one species-specific constant (*b* or *c*) between *I* and *T*, though variation in both was also possible (**Figure 2**). However, with three species, co-existence requires variation in both *b* and *c* between the species *I* and *F* (**Figure 3**). This result is easier to demonstrate when *b*_I_ = *b*_F_; in that case, the payoff difference ***P***_***I***_ – ***P***_***F***_ = *c*_F_ **−** *c*_I_ becomes independent of frequency, guaranteeing the extinction of one of the species. On the other hand, if *c*_I_ =*c*_F_, this leads to a zero equilibrium frequency for *T* (Appendix B, equation (14)), making this case impossible. Specifically, the variation in *b, c* between the two species must be a relative specialization: if *I* is superior to *F* in both parameters, then ***P***_***I***_ > ***P***_***F***_ will always hold. Therefore, *I* and *F* must show relative specialization in nutrient utilization, like the copio/oligotrophic life histories described previously. This emerging pattern suggests stable co-existence with more than three species may require payoffs with more than two parameters relating nutrients to growth: such additional parameters can be incorporated *e*.*g*., by considering aforementioned non-linear payoffs, or by considering other interspecies relationships, such as the exchange of additional metabolites^18,40^.

**Figure 3.**
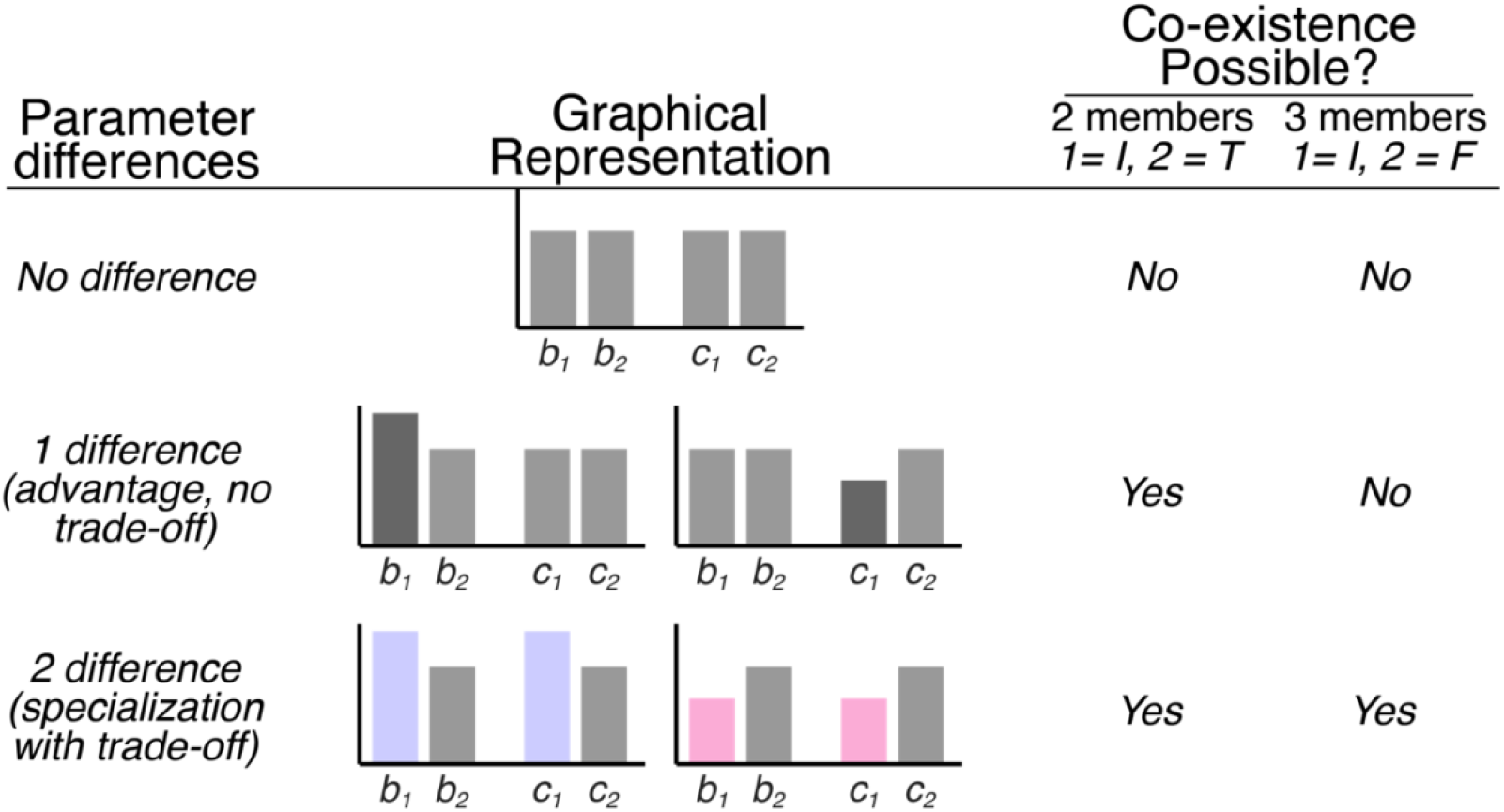
Visual summary of how stable co-existence depends on differences in species-specific parameters *b,c*, between key species in model consortia. Species subscripts 1 and 2 refer to different species in the two model consortia studied. In the two-member consortia, identical *b* and *c* parameters cannot stabilize co-existence but *I*’s dominance in a single parameter (higher *b* or lower *c*) is sufficient for stabilization. In the three-membered consortia, *I* must be specialized (better *b* and worse *c*, or vice versa) relative to *F* for stability.

For co-existence, *I* and *F* must trade-off their growth advantages. In other words, either *(c*_*F*_ >*c*_*I*_, *b*_*F*_ >*b*_*I*_ ) or *(c*_*F*_ <*c*_*I*_, *b*_*F*_ <*b*_*I*_ ) is necessary. Interestingly, we further discover that only the former condition, namely an oligotrophic *I* relative to *F*, leads to a stable co-existence equilibrium, whereas the latter condition (copiotrophic *I*) is unstable, resulting in the extinction of either *I* or *F*, depending on their initial frequencies. This is shown rigorously in Appendix B, but we emphasize that the oligotrophic *I* requirement for stability has a straightforward biological interpretation, illustrated in **Figure 4** along with an example trajectory of a stable three-membered consortium. This interpretation rests on the idea that an increase in the frequency of one species leads to an increase in the relative fitness of a single other species, allowing all species to increase in frequency from rarity in a non-transitive manner (**Figure 4A**). For instance, *T* obtains its highest relative fitness when *I* is most frequent, *i*.*e*. when intermediates are produced at the highest rates, since only *T* can directly benefit from them. As *T* becomes more frequent, nutrient concentrations increase, giving the copiotrophic *F* a growth advantage. However, as *F* grows, it displaces *I* and *T* resulting in slowed biodegradation; this in turn favors the oligotrophic *I*, which survives best in the absence of the nutrient (**Figure 4B**). If instead, *I* was copiotrophic and *F* oligotrophic, the growth of *T* would lead to further growth of *I*, creating a positive feedback loop between *I* and *T* that would exclude *F*. This explains why this regime does not result in stable co-existence with *T*. The model prediction for stability also requires the system to avoid becoming extinct during low nutrient conditions, which happens during *I*’s advantage. That is, as *I* becomes relatively more common, all species may be dying at different rates. Thus *T* must eventually obtain an advantage before any species is eliminated from the consortium.

**Figure 4.**
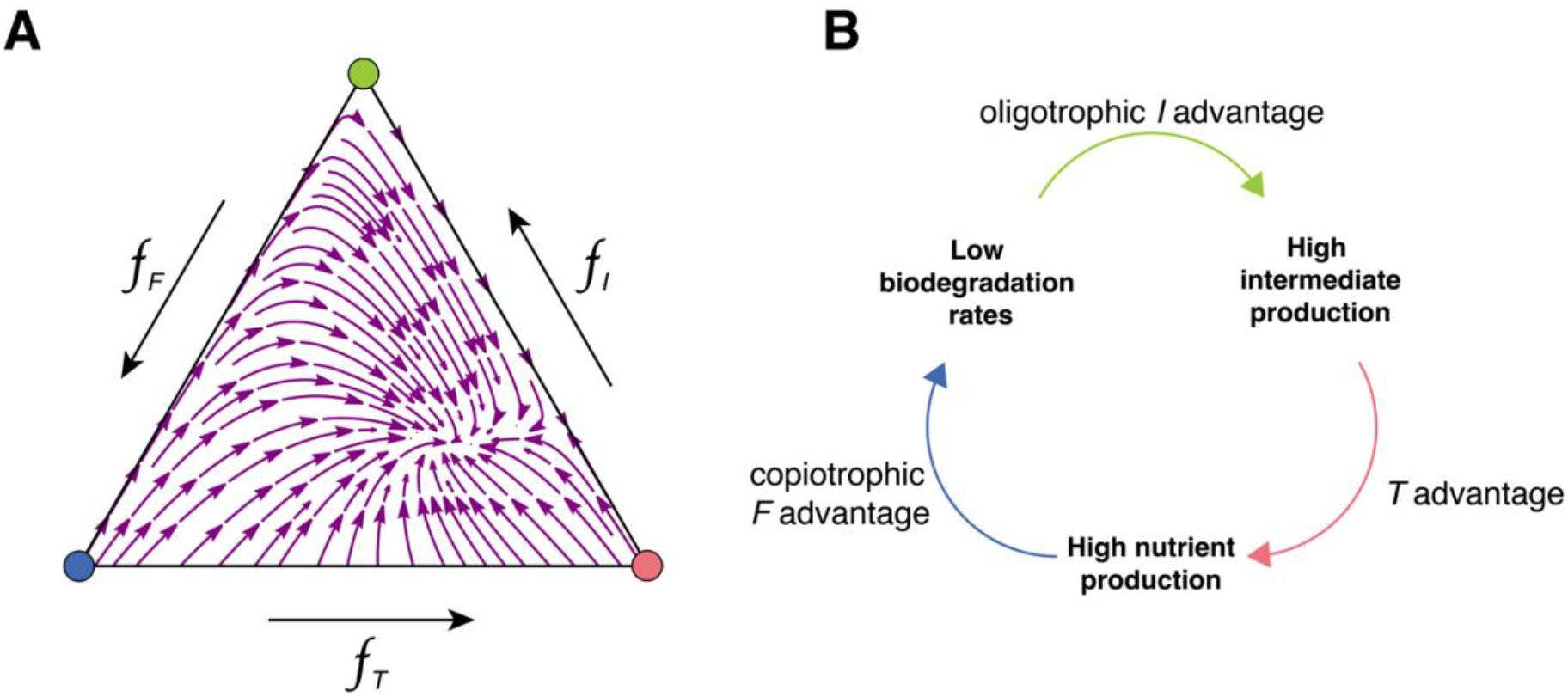
(A) Ternary simplex plot showing trajectories of a stable three species consortium. (B) Visual schematic describing how stability between the three species is manifested by each species creating favorable conditions for the other; favorable meaning higher relative fitness than both the others. The model requires that *I* creates favorable conditions for *T* by generating nutrients; *T* generating favorable conditions for *F* by generating nutrients; and *F* generating favorable conditions for *I* by consuming the nutrients and displacing both *I* and *T*. The system experiences an extinction threat during *I* ‘s advantage phase since all species may be dying at different rates. Parameter values used for simulations of (A) are *b*_*I*_ *= 4; b*_*T*_ *= 11/4; b*_*F*_ *= 8; c*_*I*_ *= 1/2; c*_*T*_ *= 1; c*_*F*_ *= 1; r*_*I*_ *= 21/22; r*_*T*_ *= 1*.

## Discussion

In order to obtain an ecologically stable EBC in laboratory enrichments, growth rates of multiple species on a single limiting nutrient must be balanced. Previous theoretical studies into the formation of interspecific groups (containing multiple species) are scarce, as genetic relatedness cannot be invoked to explain the coincidence of all species’ fitness interests^41^. Our study suggests that if the existence of diverse life histories can indeed balance asymmetric benefits and free-riding, then the fact that EBC are composed of genetically unrelated taxa, to some degree, facilitates their stable formation. This conclusion must obviously be handled with care: while metabolic trade-offs support coexistence, they do not inherently bias consortia toward mutual benefit over exclusion. Stability depends on “ecological matching” between members, such as an oligotrophic *I* and a copiotrophic *F* species. Without this balance, xenobiotic-degrading consortia risk instability on an ecological timescale. The extent to which ecological matching can be satisfied in natural scenarios is not demonstrable, assuming that unsuccessful enrichments are not reported in the literature. Hardly any EBC enrichment experiments report the results of multiple replicate enrichments, and results are variable^42,43^. Nonetheless, the ability to enrich consortia on a variety of xenobiotics over many decades of research supports a widespread potential for ecological matching^6^.

Species like *I* seem to play a problematic role within EBC, performing steps that provide no direct benefit. Historically considered “co-metabolic”^12,18,44,45^, *I* species were thought to depend on external nutrients for stability, but they are known to coexist within EBC, feeding off excess nutrients from other members. For example, in a consortium degrading the pesticide parathion, the *I* species *Pseudomonas stutzeri* produced *para*-nitrophenol, which it could not utilize, but was metabolized by *Pseudomonas aeruginosa*^44^. Carbon from *P. aeruginosa*’s metabolites and cell lysis products supported *P. stutzeri* and two non-metabolizing species^44^. This EBC remained stable for over two years in a chemostat. Interestingly, *P. stutzeri* was noted as “slower-growing”; this may seem surprising in the face of its need to compete with other non-metabolizers for the carbon released by *P. aeruginosa*. However, our model aligns with these observations, predicting further that the slow-growth should trade off for increased survival at low-nutrient concentrations.

### Ecological implications for metabolic evolution

Ecological matching between species appears restrictive of an already limited probability, namely, discovering all necessary reactions for a new pathway from promiscuous enzymes within a community. This probability need not, however, be so slight. For example, microbial enzymes for degrading parathion have evolved independently across different protein folds^46^, and even unrelated enzymes can show some hydrolysis activity towards similar pesticides like paraoxon (*e*.*g*., some antibiotic-inactivating enzymes)^47–49^. It is well-known that some xenobiotic catabolic pathways appear to evolve faster than others, such as the rapid microbial adaptation to diphenamid^50^, versus the slower evolution of atrazine degradation, showing a sudden increase in catabolic rates after half a century of slow breakdown^15^. This suggests certain reactions are more widely present as latent promiscuous activities.

If reactions like parathion hydrolysis arise frequently in microbial communities, multiple species may form a consortium. In this case, metabolic pathway evolution is selection-limited, not mutation-limited—standing variation provides promiscuous enzymes, but the limiting step is finding the right ecological background. This contrasts a typical assumption in studies of enzyme functional evolution, which regard promiscuous activities as rare “innovations” and the limiting step in metabolic evolution^51,52^. Notably, recent studies of pathogen evolution support a similar selection-limited process ^53,54^.

### On the issue of within-species free-riding

Free-riding typically refers to a within-species phenomenon of losing a cooperative trait—*e*.*g*. generating nutrients from a xenobiotic—by mutation, allowing mutants to exploit producers^55^. Between-species mutualisms, such as those between *I* and *T* species, are doubly vulnerable to free-riding, as free-riders can arise from either species^23,56^. Functional loss of either metabolic genes in *I* or *T* should result in non-metabolizing species that are otherwise unchanged in their *b* and *c* parameters.

To resolve this vulnerability for EBC, within the simplified framework of our model, we propose an antagonistic pleiotropic link between enzymatic function and growth parameters, such that functional loss incurs a cost on the organism. This is supported by the theory of enzyme promiscuity, as new promiscuous activities often arise from variation in native enzymes, which confer a “primary” function for the species. The strength of antagonistic pleiotropy to maintaining cooperation is generally contested^57–61^. However, pleiotropic linkage is a natural consequence of sourcing new reactions from standing promiscuous variation. Furthermore, the idea that functional loss results in a viable and exploitative free-riding mutant is itself only partially predictive^62–70^. An unexplored but interesting question in this regard is if mutants with lower gene expression, conferring a cost in the reduction of primary activity, but benefit due to free-riding, can invade an EBC such as the one described here. Newfound pleiotropy can thus lead to a new optimal level of the expression level of an enzyme.

### Outstanding questions

A few key ecological phenomena remain unaddressed by the model. One is the fate of the many species co-existing at the start of the enrichment, most of which go extinct before the final stable consortium emerges^17,18,71^. This initial loss likely reflects the inability of most species to survive under laboratory conditions (uncultivability)^72,73^. A model of microbial competition has shown a similar pattern of initial extinction followed by the stabilization of a small consortium, but the connection to EBC remains unstudied^74^. Another unresolved issue stems from high-throughput sequencing, which challenged the taxonomic conclusions of early EBC studies. While early studies identified a few species morphologically, later sequencing revealed that each “species” was an assemblage of strains, which can vary over time, though community function (e.g., xenobiotic degradation) remained stable^75,76^. The link between taxonomic variation and stable function remains controversial^76–78^.

## Acknowledgements

We thank Javad Meghrazi, Karol Buda, Kimberly Wong and Samuel Munn about continual feedback through the development of the project. We also thank Ming Liu and Stuart West for feedback on the completed manuscript.

## Appendix

### Appendix A: More explicit modelling of a nutrient generation flux

The payoff functions described above may be supplanted for payoffs with more realism afforded to the effects of rate on the overall metabolic flux through a mutli-step pathway. To this end, an approximation to such a flux *F* was derived in the seminal work of Kacser and Burns^39^. They showed for a pathway composed of *M* individual enzymes with rates *E*_*1*_,*…,E*_*M*_ that the equation for the flux is of the form

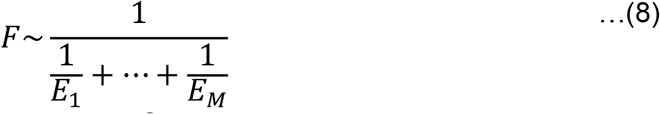

The rate terms here are composite. Specifically, equation (8) was analytically shown to hold for the flux through a pathway of substrate-unsaturated enzymes: in this case, each enzyme’s reaction rate *v* is well-approximated as linearly increasing with substrate concentration [*S*] such that 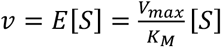, where *V*_max_ is the maximal velocity attainable under a constant enzyme concentration in the cell, and *K*_*M*_ is the half-saturation or so-called Michaelis constant^39^. Subsequent work and computer simulations showed that allowing enzyme production to vary, as well as allowing for saturated enzymes, does not significantly detract from this result, such that using equation (8) remains a good approximation^79,80^.

Following equation (8) we re-cast the metabolite concentration assumption 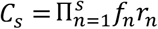 instead as 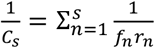. The payoffs (equations (3),(4) and (7)) then become

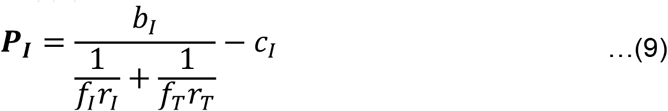

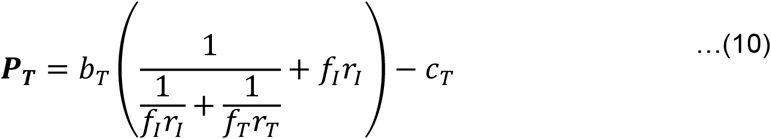

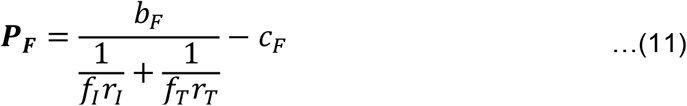

In the following paragraphs we present an abridged analysis of the *I,T* and *I,T,F* consortia, highlighting interesting modifications to the results but otherwise showing the conclusions in the main text hold. This model is hereafter referred to as the “flux” model and the model in the main text will be referred to as the “simplified” model.

For the *I,T* pair alone, the copiotrophic equilibrium conditions become stricter in the flux model, in particular β > 2. Additional conditions include *b*_T_*r*_*I*_(β **−** 2) > **−**κ, arranged to show similarity to equation (5), as well as a new condition:

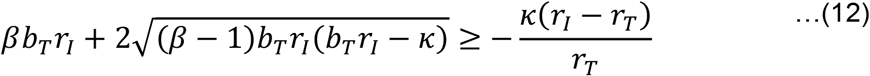

We compared the size of the parameter space allowing for stable-coexistence using these new conditions against the ones in the main text. Relative to this flux model, we find the simplified model switches from underestimating to overestimating the size of the parameter space as *r*_*T*_ gets larger; increasing *r*_*I*_ is sometimes sufficient but not necessary for this qualitative switch. The qualitative shape of the region circumscribed by these inequalities is the same and they mostly overlap except at the boundaries of small β and highly negative κ.

The *I,T* pair co-existing with an oligotrophic *I* is stable under similar conditions between the two models, assuming (6) as before (see main text). Furthermore, in the three-species consortium we find the same conclusions regarding stable co-existence as the ones described in the main text, *i*.*e*. requiring *I* to be oligotrophic relative to *F*.

### Appendix B: Why an oligotrophic I species is necessary for stable co-existence

The following argument utilizes the difference between the payoffs *D* = ***P***_***I***_ **− *P***_***F***_ ; this quantity does not represent differences in fitness but can be useful to highlight certain dynamical features, as *e*.*g*., in the main text it was used to show that co-existence is impossible when *b*_I_ =*b*_T_ =*b*_F_. Firstly, note that a single fixed point exists that contains all three species. Consider a small perturbation 1 ≫ ϵ > 0 from equilibrium frequencies such that *f*_*I*_ slightly increases and *f*_*F*_ slightly decreases above the equilibrium frequencies 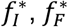. Following this perturbation, we have that

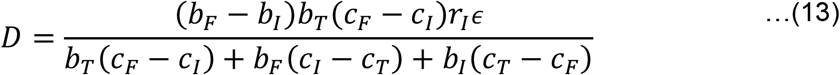

For the fixed point to be stable, we must have that *D*[ϵ] < 0. The numerator of (13) is positive regardless of whether *I* is copiotrophic or oligotrophic. This means that the denominator must be negative. However, note that the equilibrium frequency of the *T* species is

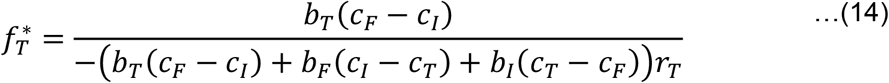

If we assume that *I* is copiotrophic, then *c*_I_ > *c*_F_, making the numerator negative; thus the denominator must be negative as well. However, the term inside the denominator parenthesis in (14) is identical to the denominator of (13), which we established must be negative. Therefore, the denominator of (14) as a whole cannot be negative. We therefore encounter a contradiction. Assuming an oligotrophic *I*, on the other hand, does not lead to such a contradiction.

