## Supplementary Information for "Stability of multi-species consortia during microbial metabolic evolution"

| Symbol | Meaning |
|---|---|
| $f_X$ | Frequency of species X in the population |
| $b_X$ | Parameter for the slope |
| $c_X$ | The baseline payoff of species X in the absence of nutrient |
| $r_X$ | Rate by which species X makes available the chemical it produces in the metabolic sequence |
| $\Phi$ | Average fitness |
| $\beta$ | The relative specialization of species I ' s b parameter relaitve to T |
| $\kappa$ | The relative specialization of species I ' s c parameter relaitve to T |

NOTE: The table above and main text utilize subscripts to indicate the species to which the parameter is attributed. In the code accompanying this supplementary file, however, we write the parameters without subscripts, since the letter "I" used to indicate one of the species is a protected symbol Mathematica used to denote the imaginary unit. Therefore the symbol $b_X$ becomes bX and so on.

#### Co-existence of *I* and *T* mutualists

##### Payoff functions and solutions

The payoffs ($P_I$, $P_T$) are defined as follows (equations (3) and (4) in main text)

```
PI = bI rI fI rT fT - cI;
PT = bT rI fI (rT fT + 1) - cT;
```

Using these payoffs we construct the replicator dynamics (equation (2) in main text) to track the fate of all genotypes in the population and identify fixed points.

In[ ]:=
```
Φ = PI fI + PT fT;
dIdt = fI (PI - Φ);
dTdt = fT (PT - Φ);
```

By setting $f_T = 1 - f_I$, we obtain a single differential equation with four solutions

In[ ]:=
```
Solve[dIdt == 0 /. fT → 1 - fI, fI] // FullSimplify
```

```
{{fI → 0}, {fI → 1},
 {fI → -((rI (bT - bI rT + bT rT) + Sqrt[rI (-4 (bI - bT) (cI - cT) rT + rI (bT - bI rT + bT rT)^2)]) /
      (2 (bI - bT) rI rT))},
 {fI → (-rI (bT - bI rT + bT rT) + Sqrt[rI (-4 (bI - bT) (cI - cT) rT + rI (bT - bI rT + bT rT)^2)]) /
     (2 (bI - bT) rI rT)}}
```

The first two fixed points {0,1} are trivial, involving only one of the species. We are primarily concerned with the properties of the last two fixed points, which we will term $FP_1$ and $FP_2$. We analyze these fixed points under the Copiotrophy and oligotrophy regimes, respectively

```
(*Saving fixed points 3 and 4 above as FP1 and FP2*)
{FP1, FP2} = fI /. Part[%, {3, 4}];
```

#### Co-existence and species-specific parameters

Co-existence between $I$ and $T$ is not possible in our model assuming the same species-specific growth parameters: this leads to $T$ always going to fixation, as illustrated by a diference between the payoffs $P_I - P_T$ which is always negative

In[ ]:=
```
PI - PT /. cI → cT /. bI → bT /. fT → 1 - fI // FullSimplify
```

Out[ ]=
```
-bT fI rI
```

However, it is sufficient that $b_I > b_T$ or $c_I < c_T$ for the stable co-existence of both species. Below is code for plotting the relationship between $\frac{d f_I}{d t}$ and $f_I$, to illustrate examples of stable fixed points (co-existence of $I$ and $T$) under differences in only one of the two parameters

In[ ]:=
```
PlotParameters = {bT → 1, bI → 1, cT → 1, cI → 0, rI → 2, rT → 2};
dIdt /. fF → 0 /. fT → 1 - fI /. PlotParameters;
plot1 = Plot[%, {fI, 0, 1}, PlotLabel → Style["bI=bF, cI<cF", Black, FontSize → 15],
    ImageSize → Medium, AxesLabel → {Style["fI", Black, FontSize → 10], Style["d fI/dt", Black, FontSize → 10]}];

PlotParameters = {bT → 1, bI → 2, cT → 0, cI → 0, rI → 2, rT → 2};
dIdt /. fF → 0 /. fT → 1 - fI /. PlotParameters;
plot2 = Plot[%, {fI, 0, 1}, PlotLabel → Style["bI>bF, cI=cF", Black, FontSize → 15],
    ImageSize → Medium, AxesLabel → {Style["fI", Black, FontSize → 10], Style["d fI/dt", Black, FontSize → 10]}];
GraphicsRow[{plot1, plot2}]
```

Out[ ]=

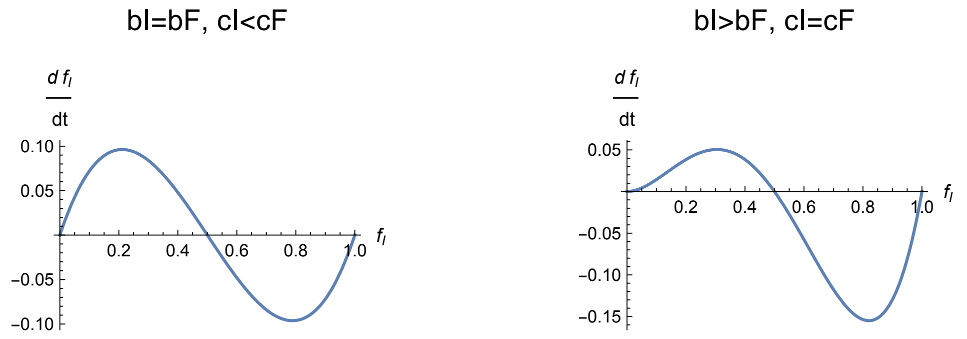


#### Co-existence under Copiotrophy

We define copiotrophy as $b_I > b_T$ and $c_I > c_T$. Using the notation $\beta = \frac{b_I}{b_T}$, $\kappa = c_T - c_I$, copiotrophy corresponds is $\beta > 1$ and $\kappa < 0$.
Note that our choice to use a ratio for $\beta$ but a difference for $\kappa$ stems from the fact that the $b$ parameters are restricted to the positive reals, whereas we allow $c$ parameter values to be negative as well as positive.

We first show that FP1 and FP2 must both exist (as frequencies between 0 and 1) under copiotrophy

In[ ]:=
```
ParameterAssumptions = bT > 0 && bI > 0 && rI > 0 && rT > 0;
Reduce[
 1 > FP2 > 0 && ! (1 > FP1 > 0) && κ < 0 && β > 1 && ParameterAssumptions /. cI → cT - κ /.
  bI → β bT]
Reduce[
 1 > FP1 > 0 && ! (1 > FP2 > 0) && κ < 0 && β > 1 && ParameterAssumptions /. cI → cT - κ /.
  bI → β bT]
```

Out[ ]= False

Out[ ]= False

To obtain the conditions described in the copiotrophy section, including equation (5), we check the conditions satisfying both fixed points being between 0 and 1
(The expression bT $\geq -\frac{4\,rT\,(-1+\beta)\,\kappa}{rI\,(1+rT-rT\,\beta)^2}$ is slightly rearranged in equation (5) in main text)

In[∘]:=
```
Reduce[1 > FP2 > 0 && 1 > FP2 > 0 && κ < 0 && β > 1 && ParameterAssumptions /. cI → cT - κ /.
    bI → β bT] // FullSimplify
```

Out[∘]= $rI > 0 \,\&\&\, \beta > 1 \,\&\&\, rT > \frac{1}{-1+\beta} \,\&\&\, \kappa < 0 \,\&\&\, bT \geq -\frac{4\,rT\,(-1+\beta)\,\kappa}{rI\,(1+rT-rT\,\beta)^2}$

In the following plot of $\frac{d f_I}{dt}$ against $f_I$, we can see that the lower fixed point is unstable

In[∘]:=
```
PlotParameters = {cI → 1/2, cT → 0, bI → 3/2, bT → 1, rT → 5, rI → 5};
dIdt /. fT → 1 - fI /. PlotParameters;
Plot[%, {fI, 0, 1},
 AxesLabel → {Style["f_I", Black, FontSize → 20], Style["d f_I / dt", Black, FontSize → 20]}]
```

Out[∘]=
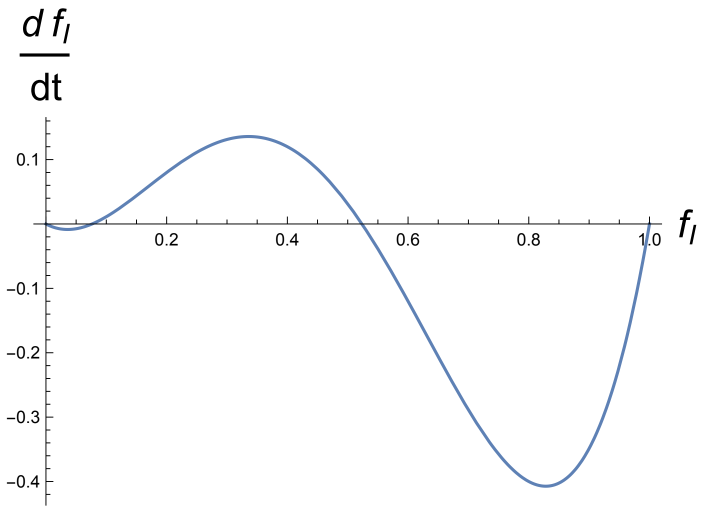


Checking the numerical values of both fixed points for this curve shows that FP1 is the unstable fixed point. We will use this when plotting both the stable and unstable

In[∘]:=
```
{FP1, FP2} /. PlotParameters // N
```

Out[∘]= {0.0763932, 0.523607}

#### Co-existence under Oligotrophy

We define oligotrophy as $b_I < b_T$ and $c_I < c_T$, equivalently $\beta < 1$ and $\kappa > 0$
The condition for $T$ to invade when rare is shown by calculating the limit of their payoff differences when $f_I$ approaches 1:

In[•]:=
```
Limit[PT - PI /. fT → 1 - fI, fI → 1] /. cI → cT - κ /. bI → β bT
```

Out[•]=
```
bT rI - κ
```

Constraining the above difference to be positive is presented in the main text as a biologically meaningful boundary for oligotrophy (see equation (6)). Briefly, the reason is that if the above quantity is negative , most of *I*'s payoff is obtained from its baseline growth rate - $c_I$, making the species unrealistically self-sufficient and making the pair not truly “co-dependent”. Therefore we require that $\kappa < b_T r_I$

Under this assumption only one of the two fixed points exists, and it is the second (stable co-existence) one

In[•]:=
```
ParameterAssumptions = bT > 0 && bI > 0 && rI > 0 && rT > 0;
Reduce[
 1 > FP1 > 0 && κ > 0 && β < 1 && ParameterAssumptions && κ < bT rI /. cI → cT - κ /. bI → β bT]
Reduce[
 1 > FP2 > 0 && κ > 0 && β < 1 && ParameterAssumptions && κ < bT rI /. cI → cT - κ /. bI → β bT]
```

Out[•]=
```
False
```

Out[•]= $rT > 0 \,\&\&\, 0 < \beta < 1 \,\&\&\, \kappa > 0 \,\&\&\, rI > 0 \,\&\&\, bT > \frac{\kappa}{rI}$

Additionally, we see that the existence of this co-existence equilibrium does not depend on additional assumptions other than the previously identified boundary $\kappa < b_T r_I$.
Below is a representative plot for co-existence with an oligotrophic *I*

In[•]:=
```
PlotParameters = {cI → 0, cT → 1/2, bI → 1, bT → 2/2, rT → 2, rI → 2};
dIdt /. fT → 1 - fI /. PlotParameters;
Plot[%, {fI, 0, 1},
 AxesLabel → {Style["f_I", Black, FontSize → 20], Style["d f_I/dt", Black, FontSize → 20]}]
```

Out[•]=
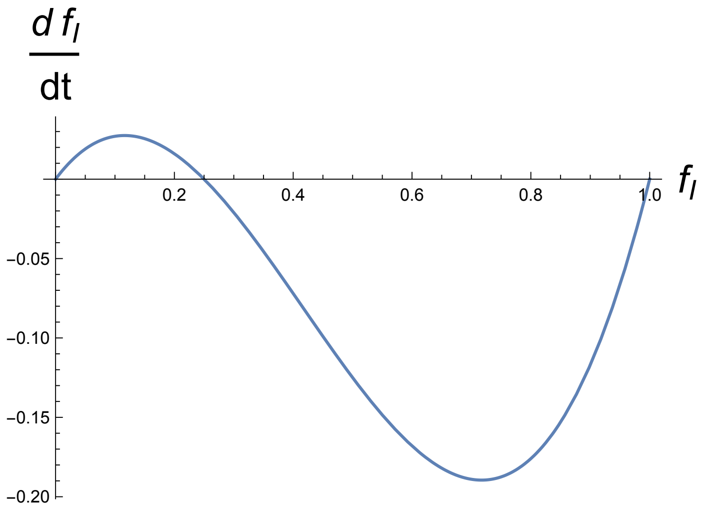

#### Figure 2

Below is a code which allows for calculating the contours used for Figure 2. The visual presentation was modified using Adobe Illustrator but the original contours were calculated here

In[•]:=
```
Show[
 ContourPlot[FP2 /. cI → cT - κ /. bI → β bT /. {rI → 5 / 4, rT → 1, bT → 1}, {β, 0, 10},
  {κ, -0.5, 1.5}, ContourShading → None, ContourStyle → Directive[Blue],
  ContourLabels → True, Contours → Flatten[{Table[i, {i, 0.1, 1.1, 0.2}]}]],
 ContourPlot[FP1 /. cI → cT - κ /. bI → β bT /. {rI → 5 / 4, rT → 1, bT → 1}, {β, 0, 10},
  {κ, -0.5, 1.5}, ContourShading → None, ContourStyle → Directive[Red, Dashed],
  ContourLabels → (Text[#3, {#1, #2}, BaseStyle → Red] &),
  Contours → Flatten[{Table[i, {i, 0, 0.3, 0.1}]}]]]
```

Out[•]=
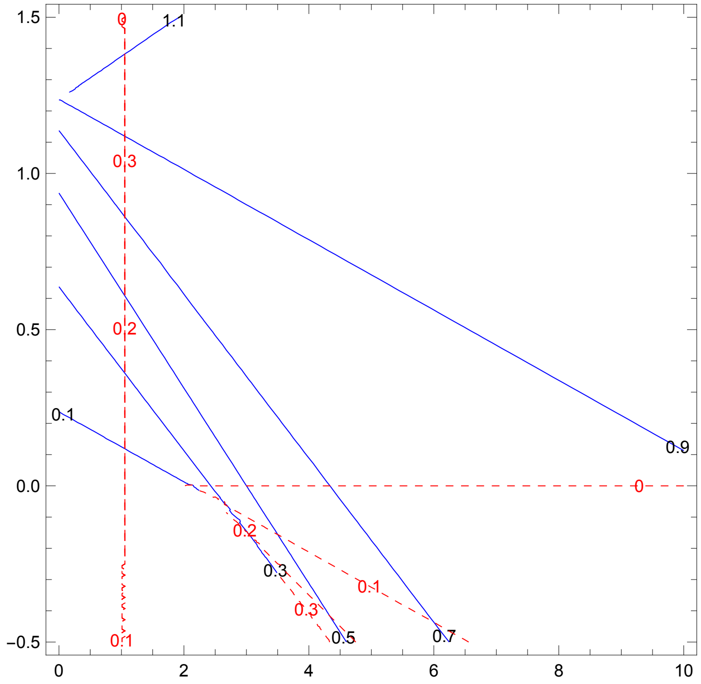


### Co-existence of *I* and *T* with free rider *F*

#### Payoff functions and solutions

The payoffs ($P_I$, $P_T$, $P_F$) are defined as follows (equations (3) (4), and (7) in main text)

In[•]:=
```
PI = bI rI fI rT fT - cI;
PT = bT rI fI (rT fT + 1) - cT;
PF = bF rI fI rT fT - cF;
```

The fixed points for this three-species systems are calculated as before, to yield:

In[ ]:=
```
Φ = PI fI + PT fT + PF fF;
dIdt = fI (PI - Φ);
dTdt = fT (PT - Φ);
dFdt = fF (PF - Φ);
Solve[dIdt == 0 && dFdt == 0 /. fT → 1 - fI - fF, {fI, fF}] // FullSimplify
```

Out[ ]=
$$\left\{\{fI \to 0, fF \to 0\}, \{fI \to 0, fF \to 1\}, \{fI \to 1, fF \to 0\}, \left\{fI \to \frac{bI\,cF - bT\,cF - bF\,cI + bT\,cI + bF\,cT - bI\,cT}{bF\,bT\,rI - bI\,bT\,rI}, fF \to 1 - \frac{bI\,cF - bT\,cF - bF\,cI + bT\,cI + bF\,cT - bI\,cT}{bF\,bT\,rI - bI\,bT\,rI} + \frac{bT\,(cF - cI)}{(bT\,(cF - cI) + bF\,(cI - cT) + bI\,(-cF + cT))\,rT}\right\}, \left\{fI \to -\left(\left(rI\,(bT - bI\,rT + bT\,rT) + \sqrt{\left(rI\,\left(-4\,(bI - bT)\,(cI - cT)\,rT + rI\,(bT - bI\,rT + bT\,rT)^2\right)\right)}\right) \Big/ (2\,(bI - bT)\,rI\,rT)\right), fF \to 0\right\}, \left\{fI \to \left(-rI\,(bT - bI\,rT + bT\,rT) + \sqrt{\left(rI\,\left(-4\,(bI - bT)\,(cI - cT)\,rT + rI\,(bT - bI\,rT + bT\,rT)^2\right)\right)}\right) \Big/ (2\,(bI - bT)\,rI\,rT), fF \to 0\right\}\right\}$$

Notably, there is only a single fixed point containing all three species. We will save these equilibrium frequencies for *I*, *F* and *T* as *Istar, Fstar* and *Tstar,* respectively:

In[ ]:=
```
{Istar, Fstar, Tstar} =
  {fI /. Part[%, 4], fF /. Part[%, 4], 1 - fI - fF /. Part[%, 4]} // FullSimplify
```

Out[ ]=
$$\left\{\frac{bI\,cF - bT\,cF - bF\,cI + bT\,cI + bF\,cT - bI\,cT}{bF\,bT\,rI - bI\,bT\,rI}, 1 - \frac{bI\,cF - bT\,cF - bF\,cI + bT\,cI + bF\,cT - bI\,cT}{bF\,bT\,rI - bI\,bT\,rI} + \frac{bT\,(cF - cI)}{(bT\,(cF - cI) + bF\,(cI - cT) + bI\,(-cF + cT))\,rT}, \frac{bT\,(cF - cI)}{(bT\,(-cF + cI) + bI\,(cF - cT) + bF\,(-cI + cT))\,rT}\right\}$$

#### Stability of three-species co-existence

We now demonstrate that stable co-existence requires differences in both the *b* and the *c* parameter (Figure 3).

Supposing that $b_I = b_F$, then the payoff difference between the two species is frequency-independent, precluding the possibility of negative frequency-dependent equilibrium

In[ ]:=
```
PI - PF /. bF → bI // FullSimplify
```

Out[ ]=
```
cF - cI
```

Next, supposing that $c_I = c_F$, we ind that this leads to an equilibrium frequency of zero for species *T*:

In[•]:=
```
Tstar /. cF → cI
```

Out[•]=
```
0
```

Therefore we conclude that differences in both *b* and *c* parameters are necessary for species co-existence, whereas with only *I* and *T* differences in only one of the two parameters was sufficient.

We now proceed to show that, when *I* is copiotrophic relative to *F* ($c_I > c_F$ and $b_I > b_F$) , co-existence is not possible in our model. For this, we first calculate the difference in the payoffs under a slight perturbation from equilibrium $\epsilon$

In[•]:=
```
PI - PF /. fT → 1 - fI - fF /. fI → Istar + ϵ /. fF → Fstar - ϵ // FullSimplify
```

Out[•]=
$$\frac{(bF - bI)\, bT\, (cF - cI)\, rI\, \epsilon}{bT\,(cF - cI) + bF\,(cI - cT) + bI\,(-cF + cT)}$$

Note that when $\epsilon$ is zero, the payoff difference is zero, since the species are then in equilibrium. Furthermore, note that this perturbation slightly increases $f_I$ relative to $f_F$; thus, for the species to return to an equilibrium, we require that this payoff difference will be negative. This reduces to the following set of inequalities

In[•]:=
```
ParameterAssumptions =
  bI > bF > 0 && bT > 0 && rI > 0 && rT > 0 && cI ∈ ℝ && cT ∈ ℝ && cF ∈ ℝ && cI > cF;
Reduce[((bF - bI) bT (cF - cI) ϵ rI)/(bT (cF - cI) + bF (cI - cT) + bI (-cF + cT)) < 0 && ϵ > 0 && ParameterAssumptions] //
 FullSimplify
```

Out[•]=
```
rT > 0 && bF > 0 && bT > 0 && ϵ > 0 && rI > 0 &&
 bF < bI && cF < cI && bT cF + bF cI + bI cT < bI cF + bT cI + bF cT
```

Note that the last inequality, $b_T\, c_F + b_F\, c_I + b_I\, c_T < b_I\, c_F + b_T\, c_I + b_F\, c_T$ requires that the denominator of the equilibrium frequency *Tstar* is positive

In[•]:=
```
Reduce[Denominator[Tstar] ≤ 0 &&
  bT cF + bF cI + bI cT < bI cF + bT cI + bF cT && ParameterAssumptions]
```

Out[•]=
```
False
```

If the denominator of *Tstar* is positive, its numerator must be positive as well; however, this would require that $c_I < c_F$, which leads to a contradiction.

In[•]:=
```
Numerator[Tstar]
```

Out[•]=
```
bT (cF - cI)
```

This contradiction can be shown directly using the Reduce function. Note that no such contradiction occurs if we instead assume *I* is oligotrophic relative to *F*.

```
In[•]:= Reduce[((bF - bI) bT (cF - cI) ϵ rI)/(bT (cF - cI) + bF (cI - cT) + bI (-cF + cT)) < 0 &&
          ϵ > 0 && 1 > Tstar > 0 && ParameterAssumptions] // FullSimplify
        OligotrophicIParameterAssumptions =
          bF > bI > 0 && bT > 0 && rI > 0 && rT > 0 && cI ∈ ℝ && cT ∈ ℝ && cF ∈ ℝ && cF > cI;
        Reduce[((bF - bI) bT (cF - cI) ϵ rI)/(bT (cF - cI) + bF (cI - cT) + bI (-cF + cT)) < 0 && ϵ > 0 &&
          1 > Tstar > 0 && OligotrophicIParameterAssumptions] // FullSimplify

Out[•]= False

Out[•]= bI > 0 && bT > 0 && ϵ > 0 && rI > 0 && bI < bF &&
         cI < cF && bT cF + bF cI + bI cT < bI cF + bT cI + bF cT &&
         bF (cI - cT) rT + bI (-cF + cT) rT + bT (cF - cI) (1 + rT) < 0
```

Beyond the conditions established above, we did not conduct a complete stability analysis of the co-existence equilibrium. However, we do provide the two eigenvalues below, describing linearized behaviour around the three species equilibrium. These eigenvalues are lengthy, but they are of the form *A(B± Sqrt[C]);* we also provide these three elements below for further analysis.

```
In[•]:= ( D[dIdt, fI]  D[dIdt, fF]
          D[dFdt, fI]  D[dFdt, fF] ) /. fT → 1 - fI - fF /. {fI → Istar, fF → Fstar} // Simplify;
        Factor[Det[% - λ IdentityMatrix[2]]] // Simplify;
        {eigen1, eigen2} = λ /. Solve[% == 0, λ] // Simplify

{-((bF^2 bT^2 cF^2 rI - bF bI bT^2 cF^2 rI - 2 bF^2 bT^2 cF cI rI + 2 bF bI bT^2 cF cI rI + bF^2 bT^2 cI^2 rI - bF bI bT^2 cI^2 rI - bF^2 bT^2 cF cT rI + 2 bF bI bT^2 cF cT rI - bI^2 bT^2 cF cT rI + bF^2 bT^2 cI cT rI -
    2 bF bI bT^2 cI cT rI + bI^2 bT^2 cI cT rI + bF bI^2 cF^3 rT - bI^3 cF^3 rT - 2 bF bI bT cF^3 rT + 2 bI^2 bT cF^3 rT + bF bT^2 cF^3 rT - bI bT^2 cF^3 rT - 2 bF^2 bI cF^2 cI rT + bF bI^2 cF^2 cI rT +
    bI^3 cF^2 cI rT + 2 bF^2 bT cF^2 cI rT + 2 bF bI bT cF^2 cI rT - 4 bI^2 bT cF^2 cI rT - 3 bF bT^2 cF^2 cI rT + 3 bI bT^2 cF^2 cI rT + bF^3 cF cI^2 rT + bF^2 bI cF cI^2 rT - 2 bF bI^2 cF cI^2 rT -
    4 bF^2 bT cF cI^2 rT + 2 bF bI bT cF cI^2 rT + 2 bI^2 bT cF cI^2 rT + 3 bF bT^2 cF cI^2 rT - 3 bI bT^2 cF cI^2 rT - bF^3 cI^3 rT + bF^2 bI cI^3 rT + 2 bF^2 bT cI^3 rT - 2 bF bI bT cI^3 rT - bF bT^2 cI^3 rT +
    bI bT^2 cI^3 rT + 2 bF^2 bI cF^2 cT rT - 4 bF bI^2 cF^2 cT rT + 2 bI^3 cF^2 cT rT - 2 bF^2 bT cF^2 cT rT + 4 bF bI bT cF^2 cT rT - 2 bI^2 bT cF^2 cT rT - 2 bF^3 cF cI cT rT + 2 bF^2 bI cF cI cT rT +
    2 bF bI^2 cF cI cT rT - 2 bI^3 cF cI cT rT + 4 bF^2 bT cF cI cT rT - 8 bF bI bT cF cI cT rT + 4 bI^2 bT cF cI cT rT + 2 bF^3 cI^2 cT rT - 4 bF^2 bI cI^2 cT rT + 2 bF bI^2 cI^2 cT rT - 2 bF^2 bT cI^2 cT rT +
    4 bF bI bT cI^2 cT rT - 2 bI^2 bT cI^2 cT rT + bF^3 cF cT^2 rT - 3 bF^2 bI cF cT^2 rT + 3 bF bI^2 cF cT^2 rT - bI^3 cF cT^2 rT - bF^3 cI cT^2 rT + 3 bF^2 bI cI cT^2 rT - 3 bF bI^2 cI cT^2 rT + bI^3 cI cT^2 rT -
    bF^2 bI bT cF^2 rI rT + bF bI^2 bT cF^2 rI rT + bF^2 bT^2 cF^2 rI rT - bF bI bT^2 cF^2 rI rT + bF^3 bT cF cI rI rT + bF^2 bI bT cF cI rI rT - 3 bF bI^2 bT cF cI rI rT + bI^3 bT cF cI rI rT -
    3 bF^2 bT^2 cF cI rI rT + 4 bF bI bT^2 cF cI rI rT - bI^2 bT^2 cF cI rI rT - 2 bF^3 bT cI^2 rI rT + 3 bF^2 bI bT cI^2 rI rT - bF bI^2 bT cI^2 rI rT + 2 bF^2 bT^2 cI^2 rI rT - 3 bF bI bT^2 cI^2 rI rT +
    bI^2 bT^2 cI^2 rI rT - bF^3 bT cF cT rI rT + 2 bF^2 bI bT cF cT rI rT - bF bI^2 bT cF cT rI rT + 2 bF^3 bT cI cT rI rT - 5 bF^2 bI bT cI cT rI rT + 4 bF bI^2 bT cI cT rI rT - bI^3 bT cI cT rI rT -
    √((bF - bI)^2 (4 bT (cF - cI) (-bI cF + bF cI) (bT (cF - cI) + bF (cI - cT) + bI (-cF + cT)) rI rT (bT^2 (cF - cI)^2 rT + bI^2 (cF - cT) (cF - cT + bT rI) rT + bF^2 (cI - cT) (cI - cT + bT rI) rT -
          bI bT (cF - cI) (2 (cF - cT) rT + bT rI (1 + rT)) + bF (2 bI (cF - cT) (-cI + cT) rT - bT (2 cI^2 - 2 cI cT + bI cI rI - 2 bI cT rI + cF (-2 cI + 2 cT + bI rI)) rT + bT^2 (cF - cI) rI (1 + rT))) +
        (bT^2 (cF - cI)^3 rT + bI^2 (cF - cT) (cF^2 - cF (cI + cT) + cI (cT - bT rI)) rT + bF^2 (cI - cT) (cI (-cI + cT - 2 bT rI) + cF (cI - cT + bT rI)) rT + bF (-2 bI (cF - cI) (cF - cT) (cI - cT) rT +
            bT (cF^2 (2 cI - 2 cT - bI rI) + cI (2 cI^2 - 2 cI cT + bI cI rI - 3 bI cT rI) + cF (-4 cI^2 + 4 cI cT + 2 bI cI rI + bI cT rI)) rT + bT^2 (cF - cI) rI (cF - cI - cT + cF rT - 2 cI rT)) + bI
          bT (cF - cI) (-2 (cF - cI) (cF - cT) rT + bT rI (cT + cI rT)))^2))) / (2 (bF - bI)^2 bT (bT (cF - cI) + bF (cI - cT) + bI (-cF + cT)) rI rT)),
 -((bF^2 bT^2 cF^2 rI - bF bI bT^2 cF^2 rI - 2 bF^2 bT^2 cF cI rI + 2 bF bI bT^2 cF cI rI + bF^2 bT^2 cI^2 rI - bF bI bT^2 cI^2 rI - bF^2 bT^2 cF cT rI + 2 bF bI bT^2 cF cT rI - bI^2 bT^2 cF cT rI +
    bF^2 bT^2 cI cT rI - 2 bF bI bT^2 cI cT rI + bI^2 bT^2 cI cT rI + bF bI^2 cF^3 rT - bI^3 cF^3 rT - 2 bF bI bT cF^3 rT + 2 bI^2 bT cF^3 rT + bF bT^2 cF^3 rT - bI bT^2 cF^3 rT -
    2 bF^2 bI cF^2 cI rT + bF bI^2 cF^2 cI rT + bI^3 cF^2 cI rT + 2 bF^2 bT cF^2 cI rT + 2 bF bI bT cF^2 cI rT - 4 bI^2 bT cF^2 cI rT - 3 bF bT^2 cF^2 cI rT + 3 bI bT^2 cF^2 cI rT +
    bF^3 cF cI^2 rT + bF^2 bI cF cI^2 rT - 2 bF bI^2 cF cI^2 rT - 4 bF^2 bT cF cI^2 rT + 2 bF bI bT cF cI^2 rT + 2 bI^2 bT cF cI^2 rT + 3 bF bT^2 cF cI^2 rT - 3 bI bT^2 cF cI^2 rT -
    bF^3 cI^3 rT + bF^2 bI cI^3 rT + 2 bF^2 bT cI^3 rT - 2 bF bI bT cI^3 rT - bF bT^2 cI^3 rT + bI bT^2 cI^3 rT + 2 bF^2 bI cF^2 cT rT - 4 bF bI^2 cF^2 cT rT + 2 bI^3 cF^2 cT rT -
    2 bF^2 bT cF^2 cT rT + 4 bF bI bT cF^2 cT rT - 2 bI^2 bT cF^2 cT rT - 2 bF^3 cF cI cT rT + 2 bF^2 bI cF cI cT rT + 2 bF bI^2 cF cI cT rT - 2 bI^3 cF cI cT rT + 4 bF^2 bT cF cI cT rT -
    8 bF bI bT cF cI cT rT + 4 bI^2 bT cF cI cT rT + 2 bF^3 cI^2 cT rT - 4 bF^2 bI cI^2 cT rT + 2 bF bI^2 cI^2 cT rT - 2 bF^2 bT cI^2 cT rT + 4 bF bI bT cI^2 cT rT - 2 bI^2 bT cI^2 cT rT +
    bF^3 cF cT^2 rT - 3 bF^2 bI cF cT^2 rT + 3 bF bI^2 cF cT^2 rT - bI^3 cF cT^2 rT - bF^3 cI cT^2 rT + 3 bF^2 bI cI cT^2 rT - 3 bF bI^2 cI cT^2 rT + bI^3 cI cT^2 rT - bF^2 bI bT cF^2 rI rT +
    bF bI^2 bT cF^2 rI rT + bF^2 bT^2 cF^2 rI rT - bF bI bT^2 cF^2 rI rT + bF^3 bT cF cI rI rT + bF^2 bI bT cF cI rI rT - 3 bF bI^2 bT cF cI rI rT + bI^3 bT cF cI rI rT - 3 bF^2 bT^2 cF cI rI rT +
    4 bF bI bT^2 cF cI rI rT - bI^2 bT^2 cF cI rI rT - 2 bF^3 bT cI^2 rI rT + 3 bF^2 bI bT cI^2 rI rT - bF bI^2 bT cI^2 rI rT + 2 bF^2 bT^2 cI^2 rI rT - 3 bF bI bT^2 cI^2 rI rT + bI^2 bT^2 cI^2 rI rT -
    bF^3 bT cF cT rI rT + 2 bF^2 bI bT cF cT rI rT - bF bI^2 bT cF cT rI rT + 2 bF^3 bT cI cT rI rT - 5 bF^2 bI bT cI cT rI rT + 4 bF bI^2 bT cI cT rI rT - bI^3 bT cI cT rI rT +
    √((bF - bI)^2 (4 bT (cF - cI) (-bI cF + bF cI) (bT (cF - cI) + bF (cI - cT) + bI (-cF + cT)) rI rT (bT^2 (cF - cI)^2 rT + bI^2 (cF - cT) (cF - cT + bT rI) rT + bF^2 (cI - cT) (cI - cT + bT rI) rT -
          bI bT (cF - cI) (2 (cF - cT) rT + bT rI (1 + rT)) + bF (2 bI (cF - cT) (-cI + cT) rT - bT (2 cI^2 - 2 cI cT + bI cI rI - 2 bI cT rI + cF (-2 cI + 2 cT + bI rI)) rT + bT^2 (cF - cI) rI (1 + rT))) +
        (bT^2 (cF - cI)^3 rT + bI^2 (cF - cT) (cF^2 - cF (cI + cT) + cI (cT - bT rI)) rT + bF^2 (cI - cT) (cI (-cI + cT - 2 bT rI) + cF (cI - cT + bT rI)) rT + bF (-2 bI (cF - cI) (cF - cT) (cI - cT) rT +
            bT (cF^2 (2 cI - 2 cT - bI rI) + cI (2 cI^2 - 2 cI cT + bI cI rI - 3 bI cT rI) + cF (-4 cI^2 + 4 cI cT + 2 bI cI rI + bI cT rI)) rT + bT^2 (cF - cI) rI (cF - cI - cT + cF rT - 2 cI rT)) + bI
          bT (cF - cI) (-2 (cF - cI) (cF - cT) rT + bT rI (cT + cI rT)))^2))) / (2 (bF - bI)^2 bT (bT (cF - cI) + bF (cI - cT) + bI (-cF + cT)) rI rT))}
```

In[ ]:=
```
eigenA = 1/(2 (bF - bI)^2 bT (bT (cF - cI) + bF (cI - cT) + bI (-cF + cT)) rI rT);
eigenB = -((bF - bI) (bF bT^2 (cF - cI) (cF - cI - cT) rI +
        bI bT^2 (cF - cI) cT rI + bF^2 (cI - cT) ((cF - cI) (cI - cT) + bT (cF - 2 cI) rI) rT +
        (bT (-cF + cI) + bI (cF - cT)) ((cF - cI) (bT (-cF + cI) + bI (cF - cT)) - bI bT cI rI)
         rT + bF (2 (cF - cI) (cI - cT) (bT (cF - cI) + bI (-cF + cT)) +
            bT (bT (cF - 2 cI) (cF - cI) + bI (-cF^2 + 2 cF cI + cI^2 + cF cT - 3 cI cT)) rI) rT));
eigenC = ((bF - bI)^2 ((bF bT^2 (cF - cI) (cF - cI - cT) rI + bI bT^2 (cF - cI) cT rI + bF^2 (cI - cT)
             ((cF - cI) (cI - cT) + bT (cF - 2 cI) rI) rT + (bT (-cF + cI) + bI (cF - cT))
             ((cF - cI) (bT (-cF + cI) + bI (cF - cT)) - bI bT cI rI) rT +
           bF (2 (cF - cI) (cI - cT) (bT (cF - cI) + bI (-cF + cT)) + bT
                 (bT (cF - 2 cI) (cF - cI) + bI (-cF^2 + 2 cF cI + cI^2 + cF cT - 3 cI cT)) rI) rT)^2 +
         4 bT (cF - cI) (-bI cF + bF cI) (bT (cF - cI) + bF (cI - cT) + bI (-cF + cT))
          rI rT ((bF - bI) bT^2 (cF - cI) rI + (bT (-cF + cI) + bI (cF - cT) + bF (-cI + cT))
             (bF (-cI + cT) - bT (cF - cI + bF rI) + bI (cF - cT + bT rI)) rT)));
(*demonstrating that we have obtained the correct
 decomposition of the eigenvalue into its three components*)
eigen1 - (eigenA (eigenB + Sqrt[eigenC])) // PowerExpand // Simplify
eigen2 - (eigenA (eigenB - Sqrt[eigenC])) // PowerExpand // Simplify
```

Out[ ]= 0

Out[ ]= 0

#### Ternary plot for Figure 4

Below we provide the code for plotting the ternary simplex plots (2D simplex representations of all 3 species' frequencies) shown in Figure 4. The function below “FrameToEpilogS3” calculates the vector field for a given set of differential equations, and displays it over the simplex.

```
In[•]:= FrameToEpilogS3[] := Module[{myEpilog, labelPos, myFrame},
    labelPos = {{-1.15, -0.05}, {1.15, -0.05}, {0.02, 1.05}};
    myFrame = Line[{{-1, 0}, {1, 0}, {0, 1}, {-1, 0}}];
    myEpilog = {myFrame}
   ];
PlotS3Field[funcs_, vars_, col_] := Module[{thisEq, p, π, myEpilog, plotOpts},
    myEpilog = FrameToEpilogS3[]; (*draw the frame using the function above*)
    thisEq = If[Abs[p] + π > 1, (*define the equations for drawing the streamplot*)
       {0, 0},
       {funcs[[3]] - funcs[[1]], funcs[[2]]} /. {vars[[3]] → (1 - π + p) / 2, vars[[2]] → π, vars[[1]] → (1 - π - p) / 2}
      ];
    (*add colored points to the frame*)
    AppendTo[myEpilog, {{PointSize[0.05], RGBColor["Black"], Point[{0, 1}]}}];
    AppendTo[myEpilog, {{PointSize[0.05], RGBColor["Black"], Point[{-1, 0}]}}];
    AppendTo[myEpilog, {{PointSize[0.05], RGBColor["Black"], Point[{1, 0}]}}];
    (*AppendTo[myEpilog,{Text[Style["O",Large],{-0.44721359+(1-0.42229-0.44721359),0.42229}]}];*)
    (*first coordinate = p3-p1, second coordinate = p2*)
    AppendTo[myEpilog, {{PointSize[0.045], RGBColor["Orange"], Point[{0, 1}]}}];
    AppendTo[myEpilog, {PointSize[0.045], RGBColor["Blue"], Point[{-1, 0}]}];
    AppendTo[myEpilog, {{PointSize[0.045], RGBColor["Red"], Point[{1, 0}]}}];
    StreamPlot[thisEq, {p, -1.0, 1.0}, {π, 0.0, 1.0}, (*draw the streamplot on the 2-simplex*)
     AspectRatio → √3 / 2,
     Frame → False,
     StreamPoints → Fine,
     StreamScale → 0.1,
     StreamColorFunction → (col &),
     StreamStyle → Medium,
     Epilog → myEpilog (*draw the stream plot on the 2-simplex*)]];
```

We use the Mathematica function “FindInstance” to automatically generate parameters that satisfy a given set of conditions. Here we generate conditions for oligotrophic and copiotrophic *I* relative to *F* to illustrate a stable co-existence equilibrium vs. an unstable equilibrium.

To the requirements described above we additionally add specific frequencies for *I,T* and *F,* which are relatively even; this is only for ease of visualization, as with similar frequencies the fixed point is clearly visible at the centre of the simplex.

In[•]:=
```
(*This parameter set represents a oligotrophic I relative to F*)
StableCoexist = FindInstance[
    Tstar == 2/4 && Istar == 1/4 && Fstar == 1/4 && OligotrophicIParameterAssumptions &&
     cF > cI && bF > bI && bT rI > cT - cI, {bI, bF, bT, cI, cF, cT, rI, rT}] // Flatten
(*This parameter set represents an copiotrophic I relative to F*)
CopiotrophicParameterAssumptions =
  bI > bF > 0 && bT > 0 && rI > 0 && rT > 0 && cI ∈ ℝ && cT ∈ ℝ && cF ∈ ℝ && cI > cF;
UnstableCoexist = FindInstance[
    Tstar == 2/4 && Istar == 1/4 && Fstar == 1/4 && CopiotrophicParameterAssumptions &&
     cF < cI && bF < bI && bT rI > cT - cI, {bI, bF, bT, cI, cF, cT, rI, rT}] // Flatten
```

Out[•]= {bI → 1, bF → 2, bT → 1, cI → -1, cF → 0, cT → 0, rI → 4, rT → 2}

Out[•]= {bI → 4, bF → 2, bT → 1, cI → 1, cF → 0, cT → 0, rI → 2, rT → 2}

In[•]:=
```
(*Setting up arguments for the plotting function*)
funcsoligo = {dIdt, dTdt, dFdt} /. StableCoexist;
funcscopio = {dIdt, dTdt, dFdt} /. UnstableCoexist;
vars = {fI[t], fT[t], fF[t]};
args = Flatten[Thread[{fI, fT, fF} → #] & /@ {Flatten[vars]}];

(*plotting*)
plotoligo = PlotS3Field[(funcsoligo /. args), vars, Purple];
plotcopio = PlotS3Field[(funcscopio /. args), vars, Purple];

GraphicsRow[{plotoligo, plotcopio}]
```

Out[•]=
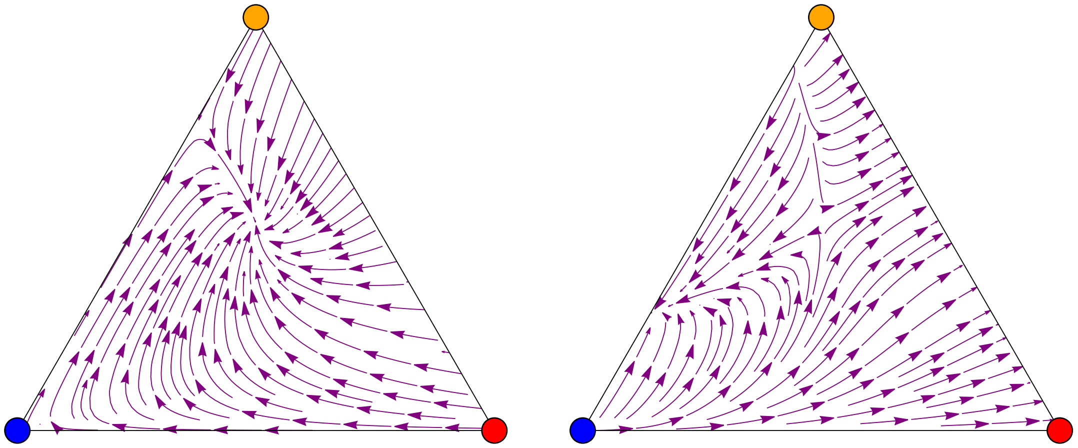

### Metabolic control-inspired payoffs

We provide an abridged analysis, reproducing the main results from the above two sections but using the more complex payoffs described in Appendix A of the main text.

The payoff functions under the assumptions of shared metabolic control of individual enzymes over the overall flux are as follows (see Appendix A in the main text for a description):

#### Payoff functions

In[ ]:=

$$PI = \frac{bI}{\frac{1}{fI\,rI} + \frac{1}{fT\,rT}} - cI;$$

$$PT = bT\left(\frac{1}{\frac{1}{fI\,rI} + \frac{1}{fT\,rT}} + fI\,rI\right) - cT;$$

$$PF = \frac{bF}{\frac{1}{fI\,rI} + \frac{1}{fT\,rT}} - cF;$$

```
Φ = PI fI + PT fT + PF fF;
dIdt = fI (PI - Φ);
dTdt = fT (PT - Φ);
dFdt = fF (PF - Φ);
```

#### *I* and *T* mutualists without *F*

We begin by reproducing the analysis of the *I* and *T* mutualists in the absence of *F*

In[ ]:=
```
Solve[dIdt == 0 /. fF → 0 /. fT → 1 - fI, fI] // FullSimplify
{FP1, FP2} = fI /. Part[%, {3, 4}];
```

Out[ ]=

$$\{\{fI \to 0\}, \{fI \to 1\}, \{fI \to -\left(\left((cI - cT)\,rI + (-cI + cT - bI\,rI + 2\,bT\,rI)\,rT + \sqrt{\left((cI - cT)^2\,rI^2 - 2\,(cI - cT)\,rI\,(cI - cT + bI\,rI)\,rT + (-cI + cT + (bI - 2\,bT)\,rI)^2\,rT^2\right)}\right) \Big/ (2\,rI\,(bT\,(rI - 2\,rT) + bI\,rT))\right)\}, \{fI \to \left(cT\,(rI - rT) + (bI - 2\,bT)\,rI\,rT + cI\,(-rI + rT) + \sqrt{\left((cI - cT)^2\,rI^2 - 2\,(cI - cT)\,rI\,(cI - cT + bI\,rI)\,rT + (-cI + cT + bI\,rI - 2\,bT\,rI)^2\,rT^2\right)}\right) \Big/ (2\,rI\,(bT\,(rI - 2\,rT) + bI\,rT))\}\}$$

Both fixed points exist for copiotrophy as before, but conditions for stable copiotrophy are a fair bit more restrictive than in the simpler payoffs case

In[∘]:=
```
ParameterAssumptions = bT > 0 && bI > 0 && rI > 0 && rT > 0;
Reduce[
 1 > FP2 > 0 && ! (1 > FP1 > 0) && κ < 0 && β > 1 && ParameterAssumptions /. cI → cT - κ /.
  bI → β bT]
Reduce[
 1 > FP1 > 0 && ! (1 > FP2 > 0) && κ < 0 && β > 1 && ParameterAssumptions /. cI → cT - κ /.
  bI → β bT]
Reduce[1 > FP2 > 0 && 1 > FP2 > 0 && κ < 0 && β > 1 && ParameterAssumptions /. cI → cT - κ /.
   bI → β bT] // FullSimplify
```

Out[∘]= False

Out[∘]= False

Out[∘]= $cT \in \mathbb{R}$ && $rI > 0$ && $rT > 0$ && $\beta > 2$ && $\kappa < 0$ &&

$$bT + \frac{\left(rT\,(-2+\beta) + rI\left(2\sqrt{\frac{(rI+rT\,(-2+\beta))\,(-1+\beta)}{rI}} + \beta\right)\right)\kappa}{rI\,rT\,(-2+\beta)^2} \geq 0$$

Oligotrophy boundary remains unchanged. The conditions themselves remain unchanged as well except for slight discrepancy when rT<rI<2 rT

In[∘]:=
```
Limit[PT - PI /. fT → 1 - fI, fI → 1] /. cI → cT - κ /. bI → β bT // Simplify
```

Out[∘]= $bT\,rI - \kappa$

In[∘]:=
```
ParameterAssumptions = bT > 0 && bI > 0 && rI > 0 && rT > 0;
Reduce[1 > FP1 > 0 && κ > 0 && 0 < β < 1 && ParameterAssumptions && κ < bT rI /. cI → cT - κ /.
  bI → β bT]
Reduce[1 > FP2 > 0 && κ > 0 && 0 < β < 1 && ParameterAssumptions && κ < bT rI /. cI → cT - κ /.
   bI → β bT] // FullSimplify
```

Out[∘]= False

Out[∘]= $cT \in \mathbb{R}$ && $bT > \frac{\kappa}{rI}$ && $rT > 0$ && $\kappa > 0$ && $\Big((0 < \beta < 1$ && $(0 < rI \leq rT \;||\; rI \geq 2\,rT))\;||$

$\Big(rT < rI < 2\,rT$ && $\Big(\Big(\beta > 0$ && $\frac{rI}{rT} + \beta < 2\Big)\;||\;\Big(\frac{rI}{rT} + \beta > 2$ && $\beta < 1\Big)\Big)\Big)\Big)$

The contour plot given the new payoffs are largely unchanged:

In[ ]:=
```
Show[
 ContourPlot[FP2 /. cI → cT - κ /. bI → β bT /. {rI → 5 / 4, rT → 1, bT → 1, cT → 0},
  {β, 0, 10}, {κ, -0.5, 1.5}, ContourShading → None, ContourStyle → Directive[Blue],
  ContourLabels → True, Contours → Flatten[{Table[i, {i, 0.1, 1.1, 0.2}]}]],
 ContourPlot[FP1 /. cI → cT - κ /. bI → β bT /. {rI → 5 / 4, rT → 1, bT → 1, cT → 0},
  {β, 0, 10}, {κ, -0.5, 1.5}, ContourShading → None,
  ContourStyle → Directive[Red, Dashed],
  ContourLabels → (Text[#3, {#1, #2}, BaseStyle → Red] &),
  Contours → Flatten[{Table[i, {i, 0, 0.3, 0.1}]}]]]
```

Out[ ]=

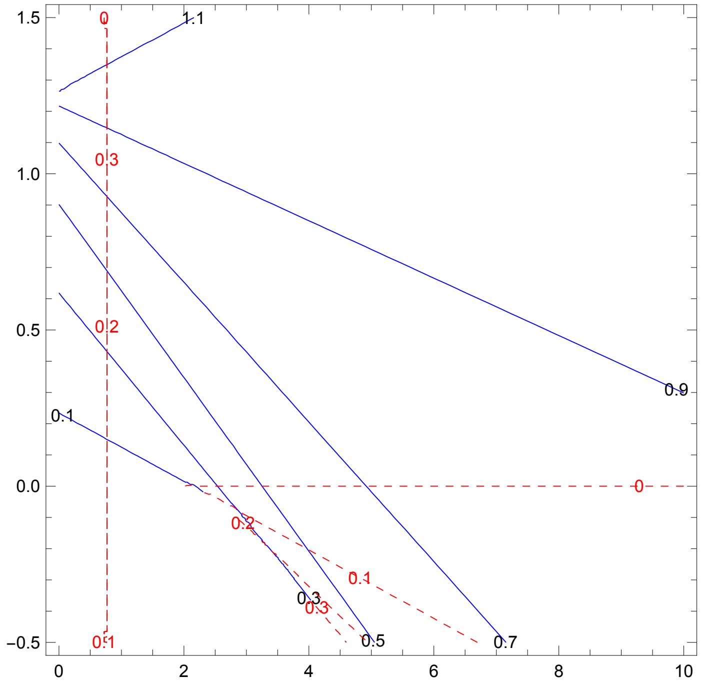


#### All three species

For three species co-existence, we find that the equilibria are similar to the ones obtained with the simpler payoffs (Istar is identical, but Fstar is not).

In[]:=
```
Solve[dIdt == 0 && dFdt == 0 /. fT → 1 - fI - fF, {fI, fF}] // FullSimplify
{Istar, Fstar, Tstar} =
  {fI /. Part[%, 3], fF /. Part[%, 3], 1 - fI - fF /. Part[%, 3]} // FullSimplify;
```

Out[]=

$$\left\{\{fI \to 0, fF \to 0\}, \{fI \to 1, fF \to 0\}, \left\{fI \to \frac{bI\,cF - bT\,cF - bF\,cI + bT\,cI + bF\,cT - bI\,cT}{bF\,bT\,rI - bI\,bT\,rI},\right.\right.$$
$$fF \to \frac{1}{bF - bI}\left(bF - bI + \frac{bT\,(cF - cI) + bF\,(cI - cT) + bI\,(-cF + cT)}{bT\,rI} + \frac{(cF - cI)\,(bT\,(cF - cI) + bF\,(cI - cT) + bI\,(-cF + cT))}{(2\,bT\,(-cF + cI) + bI\,(cF - cT) + bF\,(-cI + cT))\,rT}\right)\Bigg\},$$
$$\Big\{fI \to \Big(cT\,(rI - rT) + (bI - 2\,bT)\,rI\,rT + cI\,(-rI + rT) + \sqrt{\left((cI - cT)^2\,rI^2 - 2\,(cI - cT)\,rI\,(cI - cT + bI\,rI)\,rT + (-cI + cT + bI\,rI - 2\,bT\,rI)^2\,rT^2\right)}\Big) \Big/$$
$$(2\,rI\,(bT\,(rI - 2\,rT) + bI\,rT)),\ fF \to 0\Big\},\ \Big\{fI \to -\Big(\Big((cI - cT)\,rI + (-cI + cT - bI\,rI + 2\,bT\,rI)\,rT + \sqrt{\left((cI - cT)^2\,rI^2 - 2\,(cI - cT)\,rI\,(cI - cT + bI\,rI)\,rT + (-cI + cT + (bI - 2\,bT)\,rI)^2\,rT^2\right)}\Big) \Big/ (2\,rI\,(bT\,(rI - 2\,rT) + bI\,rT))\Big),\ fF \to 0\Big\}\Big\}$$

Requirements for differences in both *b* and *c* parameters still hold

In[]:=
```
PI - PF /. bF → bI // FullSimplify
Tstar /. cF → cI
```

Out[]= $cF - cI$

Out[]= $0$

Results of the perturbation analysis using $\epsilon$ still hold as well; however, it is now necessary to also include the assumption that 1>Istar>0 in the inequality reduction to obtain the contradiction

In[]:=
```
PI - PF /. fT → 1 - fI - fF /. fI → Istar + ϵ /. fF → Fstar - ϵ // FullSimplify
```

Out[]=

$$\left((bF - bI)\,bT^2\,(cF - cI)^2\,rI\,\epsilon\right) \Big/ \left(-(bT\,(-cF + cI) + bI\,(cF - cT) + bF\,(-cI + cT))^2 + (bF - bI)\,bT\,(2\,bT\,(cF - cI) + bF\,(cI - cT) + bI\,(-cF + cT))\,rI\,\epsilon\right)$$

In[ ]:=
```
CopiotrophicParameterAssumptions =
  bI > bF > 0 && bT > 0 && rI > 0 && rT > 0 && cI ∈ ℝ && cT ∈ ℝ && cF ∈ ℝ && cI > cF;
Reduce[((bF - bI) bT^2 (cF - cI)^2 rI ϵ) / (- (bT (-cF + cI) + bI (cF - cT) + bF (-cI + cT))^2 +
      (bF - bI) bT (2 bT (cF - cI) + bF (cI - cT) + bI (-cF + cT)) rI ϵ) < 0 &&
  ϵ > 0 && 1 > Tstar > 0 && 1 > Istar > 0 && CopiotrophicParameterAssumptions]
OligotrophicIParameterAssumptions =
  bF > bI > 0 && bT > 0 && rI > 0 && rT > 0 && cI ∈ ℝ && cT ∈ ℝ && cF ∈ ℝ && cF > cI;
Reduce[((bF - bI) bT^2 (cF - cI)^2 rI ϵ) / (- (bT (-cF + cI) + bI (cF - cT) + bF (-cI + cT))^2 +
       (bF - bI) bT (2 bT (cF - cI) + bF (cI - cT) + bI (-cF + cT)) rI ϵ) < 0 && ϵ > 0 &&
   1 > Tstar > 0 && 1 > Istar > 0 && OligotrophicIParameterAssumptions] // FullSimplify
```

Out[ ]= False

Out[ ]=
$$bI > 0 \,\&\&\, bF > bI \,\&\&\, cF > cI \,\&\&\, bT > 0 \,\&\&\, cT > \frac{-bI\,cF + 2\,bT\,(cF - cI) + bF\,cI}{bF - bI} \,\&\&$$
$$rT > \frac{(cF - cI)\,(bT\,(cF - cI) + bF\,(cI - cT) + bI\,(-cF + cT))}{(bF - bI)\,(2\,bT\,(cF - cI) + bF\,(cI - cT) + bI\,(-cF + cT))} \,\&\&$$
$$rI > \frac{bI\,cF - bT\,cF - bF\,cI + bT\,cI + bF\,cT - bI\,cT}{bF\,bT - bI\,bT} \,\&\&\, \epsilon > 0$$
